# Measurement precision bounds on aberrated single molecule emission patterns

**DOI:** 10.1101/2023.11.30.569462

**Authors:** Li Fang, Fang Huang

## Abstract

Single-Molecule Localization Microscopy (SMLM) has revolutionized the study of biological phenomena by providing exquisite nanoscale spatial resolution. However, optical aberrations induced by sample and system imperfections distort the single molecule emission patterns (i.e. PSFs), leading to reduced precision and resolution of SMLM, particularly in three-dimensional (3D) applications. While various methods, both analytical and instrumental, have been employed to mitigate these aberrations, a comprehensive analysis of how different types of commonly encountered aberrations affect single molecule experiments and their image formation remains missing. In this study, we addressed this gap by conducting a quantitative study of the theoretical precision limit for position and wavefront distortion measurements in the presence of aberrations. Leveraging Fisher information and Cramér-Rao lower bound (CRLB), we quantitively analyzed and compared the effects of different aberration types, including index mismatch aberrations, on localization precision in both biplane and astigmatism 3D modalities as well as 2D SMLM imaging. Furthermore, we studied the achievable wavefront estimation precision from aberrated single molecule emission patterns, a pivot step for successful adaptive optics in SMLM through thick specimens. This analysis lays a quantitative foundation for the development and application of SMLM in whole-cells, tissues and with large field of view, providing in-depth insights into the behavior of different aberration types in single molecule imaging and thus generating theoretical guidelines for developing highly efficient aberration correction strategies and enhancing the precision and reliability of 3D SMLM.

## INTRODUCTION

SMLM utilizes photo-switchable or photo-convertible fluorescent dyes or proteins to capture isolated single molecule emission events in a series of camera frames and subsequently localizes these individual probes with a precision down to a few nanometers ^1–6^. Such resolution at the molecular level together with its labeling specificity and live cell compatibility make SMLM a unique technology to reveal previously-hidden phenomena and uncover profound insights for biological and biomedical research^7–10^.

Single molecule localization, as one of the core concepts in SMLM, involves analyzing and extracting information of molecular positions within the three-dimensional cellular space from the detected emission patterns of single fluorescent probes^11–15^ (hereafter referred as point-spread function, PSF). Photons forming these emission patterns need to travel through the biological specimen, the objective lens and the rest of the microscopy instrument before forming PSFs captured by the camera. Inevitably, caused by the inhomogeneous refractive indices of intra- and extra-cellular constituents and instrument imperfections, PSFs suffer aberrations induced by the instrument and most importantly the specimen itself. Sample and system induced aberrations distort or blur the detected single molecule emission patterns and subsequently alter the amount of information carried by the emitted photons with respect to the molecular position and the underlying wave-front shape, leading to a deterioration of localization precision and achievable resolution, especially in three-dimensional (3D) SMLM^16–18^.

Both analytical and instrumental methods have been developed to alleviate the detrimental effects of aberrations: localization using aberration-considered PSF, though retrieval an experimental or an *in situ* PSF are developed to pinpoint the single emitter centers with high precision and accuracy^19–22^; instrumentally, adaptive optics^23–25^, using a wavefront modification device such as a deformable mirror, are introduced to correct aberrations based on measured or inferred wavefront shape^26–31^. Despite the numerous methods developed to numerically account and instrumentally compensate for aberrations across different scales, a comprehensive understanding of the practical influences of these aberrations on single molecule datasets remains obscure. Our knowledge in this domain is primarily rooted in hands-on experiences and empirical descriptions. How precisely can we localize single molecules when certain aberration exists? How precisely we can determine the wavefront shape based on aberrated PSFs? The answers to these questions remain unanswered due to the lack of quantitative analysis on the information content of single molecule emission patterns.

In this work, we provide a quantitative analysis on the lower bound of estimation uncertainties for both position and wavefront distortions using Fisher information and the Cramér-Rao lower bound (CRLB). Specifically, we provided the best possible localization precision in the presence of various aberration types including index mismatch aberration^20,32,33^ and the achievable precision when estimating aberration amplitude from aberrated emission patterns for both biplane and astigmatism systems. These analyses reveal the practical effects of different aberration types regardless of localization algorithms and quantify the limit of wavefront inference for adaptive optics in single molecule imaging experiments. With a more thorough understanding of the information carried by aberrated specimens, we will further the development of information enriched PSFs, innovate efficient aberration correction methods and thus push the envelope of optical nanoscopy in resolving the ultrastructures of intra- and extra-cellular constituents through tissues and small animals.

## RESULTS

### Fisher information matrix and Cramér-Rao lower bound

The key to advancing single molecule localization microscopy lies in addressing parameter estimation problems, such as accurately determining the locations of imaged single molecules and estimating wavefront distortions. The best precision of parameter estimation is fundamentally circumscribed by the information content carried by individual photons. Therefore, evaluating the optimal precision converges to quantify the information content of the single molecule emission pattern, by computing the Fisher information matrix based on physical model of light diffraction through microscope and statistical model of photon counting. Mathematically, when a probability density function *p*(***X***; ***θ***) of random variables ***X*** satisfies the regularity condition 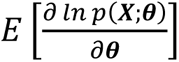 = 0 for parameter set ***θ*** = [*θ*_1_, …, *θ*_*n*_]^*T*^, the Fisher information matrix *I*(***θ***) is an *n* × *n* symmetric matrix whose *ij*-th element is defined as^34,35^:

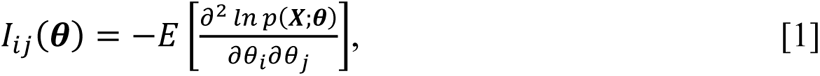

where ***θ*** is a vector of estimation parameters, ***X*** refers to the noisy data obtainable on a hypothetical detector in the case of image acquisition. In the case of single molecule imaging, ***θ*** includes the positions of the molecules, intensities, background and aberration coefficients (Zernike modes), ***X*** is the subregion of detected raw images which containing single molecules. According to Cramér-Rao inequality, the estimation variance of any unbiased estimator ***θ***_*i*_, i.e. the Cramér-Rao lower bound (CRLB), must satisfy^34^:

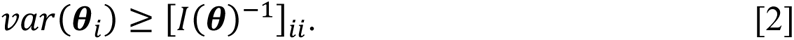

For single molecule imaging, assuming the number of photons detected in each pixel are independent random variables following a Poisson distribution and using Stirling’s approximation, the Fisher information matrix can be simplified as^36–38^:

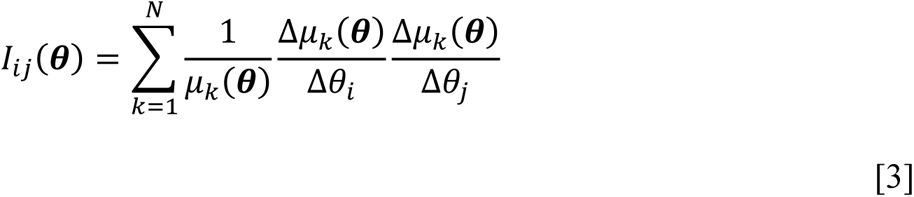

where *μ*_*k*_(***θ***) is the value of the PSF model at pixel *k*, *N* is the total number of the pixels in one fitting sub-region and ***θ*** = [*θ*_1_, …, *θ*_*n*_]^*T*^ is the parameter vector.

3D single molecule localization requires an axially unambiguous PSFs such as astigmatism modality where astigmatism wavefront distortion through a cylindrical lens is pre-introduced in the imaging system^6^ or biplane modality where a pair of PSFs from the same single molecule is detected at two axially separated planes^4^. Here, we evaluated the CRLBs for both 3D single molecule imaging systems.

For 3D SMLM systems, Fisher information matrix *I* can be calculated as the sum of the information matrices as follow^20^:

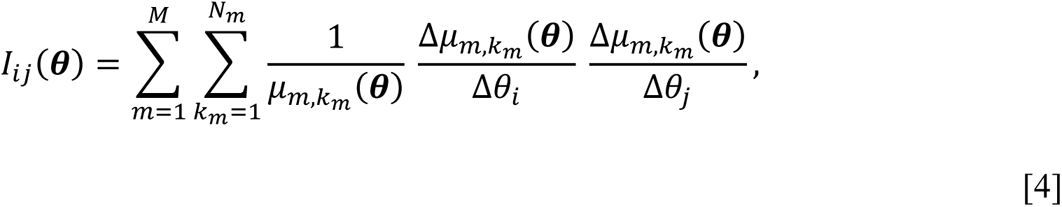

Where *M* is the number of planes, equal to 1 for astigmatism system, 2 for biplane system; *N_m_* is the pixel numbers in the *m*th plane; *μ*_*m*,*km*_(***θ***) is the value of the PSF model at pixel *k* in *m*th plan; ***θ*** is the parameter vector to be estimated.

We calculated the lower bounds of estimation precisions for 2D and 3D single molecule localization as well as wavefront estimation given an aberrated single molecule emission pattern. For calculations involving aberration estimation, the parameter set {***θ***} is given by {*x*, *y*, *z*, *I*, *bg*, *c*_5_, *c*_6_ … *c*_*i*_, … *c*_*n*_}, where (*x*, *y*, *z*) is the 3D position (unit: nm) of a single molecule, *I* is the detected photon count, *bg* is the background photon count per pixel and (*c*_5_, *c*_6_ … *c*_*i*_, … *c*_*n*_) are the amplitudes of Zernike polynomials (unit: λ/2π, Wyant order, starting from vertical astigmatism) describing a distorted wavefront shape at the pupil of an objective lens. To remove the influences of photon numbers emitted by a single molecule which improves the estimation variance with a reciprocal dependence, we quantified the precisions per photon count without background noise.

## LOCALIZATION PRECISION IN PRESENCE OF ABERRATIONS

Optical aberrations caused by instrument imperfections and inhomogeneous refractive indices of the specimen affect a single molecule imaging system through distorting and blurring the emission patterns generated from single fluorescent molecules. While distortion is capable of alternating the appearance of the emission pattern detected, not necessarily would it worsen the localization precision, particularly when an *in situ* (realistic) PSF is obtainable^19^. On the other hand, the blurring of the emission pattern, in general, lowers the position information content carried by the detected photon and therefore deteriorates localization precision. This deterioration is not recoverable using analytical methods. In the following sections, we delved into the effects of optical aberrations on localization precision.

### Lateral precision

We defined the lateral precision as the root mean square (RMS) of precisions in x- and y-direction and calculated this metric at various axial positions (−600 nm to 600 nm) with respect to the actual focus plane (z=0) in presence of different aberration types (up to 2^nd^ spherical aberration, Wyant order) and amount (up to 3 λ/2π). We calculated the lower bounds for both the biplane and astigmatism imaging modalities. The biplane distance was set at 400 nm, and the astigmatism value was fixed at 1.4 λ/2π. Under these conditions, the axial localization precision along the z-axis remained uniformly high (**Supplementary Fig.1**). We observed similar effects of aberrations on lateral localization precision in these two modalities.

In our observations, we noted that an increased magnitude of aberrations could result in both deteriorated and occasionally improved lateral precision in both the biplane and astigmatism systems (**Fig.1, Supplementary Fig.2-13**). However, these aberrations exhibited distinct behaviors that we categorized into two groups (**Fig.1a**): (1) aberrations that consistently degrade lateral precision across the axial range, such as astigmatism (vertical and diagonal, **Supplementary Fig.2-3**) (2) aberrations that deteriorate lateral precision near focus but improve it when the emitter is out of focus. The second category includes trefoil (oblique and vertical, **Supplementary Fig.7-8**), secondary astigmatism (vertical and diagonal, **Supplementary Fig.9-10**) and secondary coma (horizontal and vertical, **Supplementary Fig.11-12**). The transition point, where precision shifted from worsening to improving, typically occurred approximately 400 nm away from the focus for the biplane system and around 250 nm away for the astigmatism system. This observation suggests that, for aberration modes falling into the second category, these wavefront distortions primarily deteriorate lateral precision, with a more pronounced effect near the focus than near the boundaries of a typical axial localization range in SMLM.

**Fig. 1:**
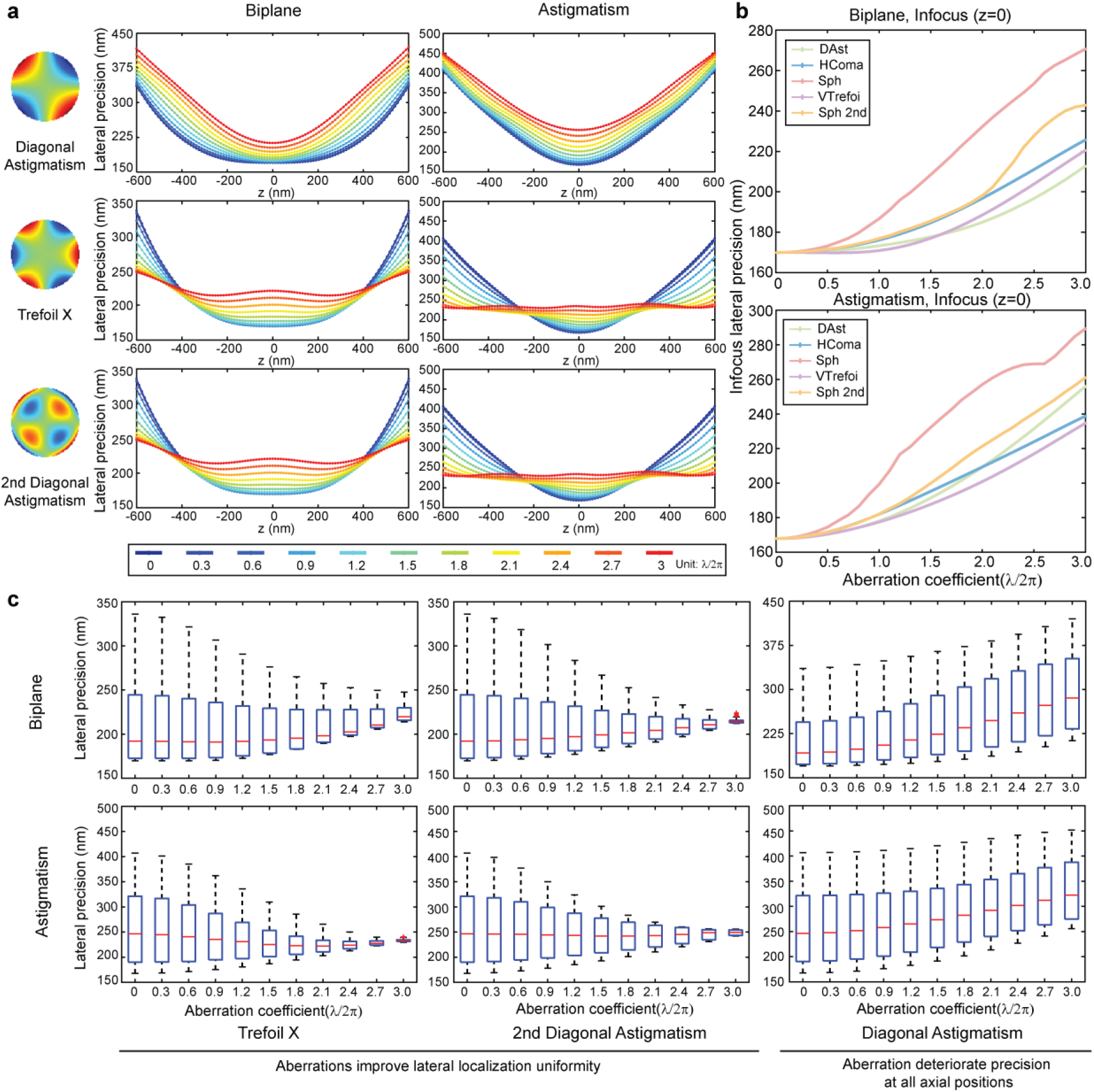
Aberration’s influences on lateral localization precision. (a) Lateral localization precision along the z-axis for diagonal astigmatism, horizontal trefoil, and secondary diagonal astigmatism in both the biplane (left) and astigmatism (right) systems. Each figure is color-coded to represent aberration coefficients ranging from 0 to 3, with a step size of 0.3, measured in units of λ/2π. (b) In-focus (at z = 0) lateral localization precision with varying aberration types and amplitudes in biplane (top) and astigmatism (bottom) systems. Abbreviations: DAst (diagonal astigmatism), HComa (horizontal coma), Sph (spherical), VTrefoil (vertical trefoil), Sph 2nd (secondary spherical). (c) Distributions of lateral localization precision within the z range of −600 nm to 600 nm in the presence of different aberration modes with varying amplitudes (ranging from 0 to 3). Each boxplot displays the median, 25th, and 75th percentiles of the data, while whiskers extend to non-outlier extreme points. Outliers are individually marked with plus signs. Simulation conditions: Wavelength (**λ**): 680 nm, intensity (*I*): 0.5 photons/plane for the biplane system and 1 photon for the astigmatism system, background (*bg*): 0, Numerical Aperture (*NA*) = 1.4, immersion medium with refractive index (*n_obj_*) = 1.52, biplane distance: 400 nm, astigmatism amplitude: 1.4 λ/2π. Lateral localization is calculated as the root mean square (RMS) of x- and y-localization precision. The region of interest (ROI) size was set to 32 × 32 pixels.

When focusing our observation on in-focus emitters (at z=0) as in the cases when planar cellular structures are imaged, we found different aberration modes deteriorate lateral precision at drastically different rates (**Fig.1b, Supplementary Fig.14**). Considering the effective focal shift of primary spherical and secondary spherical aberration, we determined the actual focal position where the PSF is mostly tightly focused by identifying the location around which the imaging range (−600 nm to 600 nm) yielded the maximum sum of peak intensities in the PSFs. This range denotes the region of highest focus, making it the optimal area for practical imaging. (**Methods, Supplementary Fig.24**). Among the tested aberration modes, primary spherical and secondary spherical aberration have the most significant impact on lateral localization at actual focal plane in both biplane and astigmatism setup (**Fig.1b**). In biplane system, the achievable lateral precision per photon count gets deteriorated from 169.9 nm to 270.7 nm (a 1.6x increase) for primary spherical aberration when the aberration amplitude increases to 3 λ/2π. Following closely, secondary spherical aberration degrades the in-focus lateral precision to 242.9 nm (from 169.9 nm, a 1.4x increase) at an amplitude of 3 λ/2π (**Supplementary Fig.14a**). In the astigmatism system, primary and secondary spherical aberrations similarly exhibit detrimental effects with a 2.7-fold and 1.6-fold precision degradation, respectively (**Supplementary Fig.14b**). In contrast, diagonal astigmatism results in the smallest effect on precision, causing only a 1.2x deterioration in the biplane setup. These observations can be intuitively understood by considering that an in-focus single molecule pattern can be blurred by primary and secondary spherical aberrations while it remains a fine focused spot in presence of astigmatism aberrations.

Aberrated wavefronts will also alter the uniformity of lateral localization precision per photon count achievable at different axial positions. Localization precision uniformity is especially important in 3D SMLM and in 2D SMLM when the cellular targets are often thicker than 500 nm, a common situation when resolving mammalian cells without total internal reflection fluorescence (TIRF) illumination^39,40^. The alteration can be observed by the amount of variation in lateral precision across the z range as the amplitude of a specific aberration mode progressively increases. The distributions of lateral precisions within a z range of −600 nm to 600 nm with different aberration amplitudes are visualized in **Fig.1c**. For each aberration type and amplitude, the interquartile range of lateral precision distribution is represented by the box length. Several types of aberrations benefit lateral localization by improving such precision uniformity. For example, the presence of primary spherical (**Supplementary Fig.6**), trefoil (oblique and vertical, **Supplementary Fig.7-8**), secondary astigmatism (vertical and diagonal, **Supplementary Fig.9-10**), secondary coma (horizontal and vertical, **Supplementary Fig.11-12**) and secondary spherical (**Supplementary Fig.13**) reduce variations of the lateral precisions at various axial positions in both biplane and astigmatism system. In contrast, vertical and diagonal astigmatism (**Supplementary Fig.2-3**) increase such variations in biplane system and have minor effects in astigmatism system. In biplane system, for diagonal astigmatism, with amplitude growing from 0 to 3 λ/2π, the interquartile range increases from 71.9 nm to 119.4 nm, demonstrating the increased lateral precision variation (**Fig.1c, Supplementary Fig.3**). On the contrary, for secondary diagonal astigmatism, the same measure decreases from 71.9 nm to 2.9 nm with an increasing aberration amplitude from 0 to 3 λ/2π, indicating a precision equalization effect of secondary astigmatism (**Supplementary Fig.10**). This strong equalization effect can also be observed in trefoil (oblique and vertical, **Supplementary Fig.7-8**), secondary spherical (**Supplementary Fig.13**) and secondary coma (horizontal and vertical, **Supplementary Fig.11-12**). These observations suggest a potential strategy of introducing specific aberrations to enhance the overall 3D resolution across the entire imaging volume for both the biplane and astigmatism imaging modalities.

### Axial precision

While the lateral position of a single molecule is determined by pin-pointing the center of its emission pattern, its axial position in commonly used astigmatism and biplane systems is extracted from the shape of the emission pattern (a.k.a. the PSF shape). Thus, the alternations of PSF shapes, induced by aberrations, change the amount of information carried by the emission pattern, heavily affecting achievable axial localization precision per photon count. As revealed in this section, these changes can be quite complex, exhibiting variations at different axial positions and in response to different aberration types and amplitudes. Importantly, while some aberration modes deteriorate axial precisions rapidly, the presence of a few commonly encountered aberration modes has minimal influences on the axial resolution. In general, the effects of aberrations on axial precision have similar trends for biplane and astigmatism system but vary in some aberrations due to the pre-introduced astigmatism in astigmatism-based 3D imaging modality.

Based on the axially dependent effects of aberrations, we group the aberration modes into 2 categories: (1) Aberrations that consistently deteriorate the axial precisions at all evaluated axial locations. This category includes coma (horizontal and vertical, **Fig.2a, Supplementary Fig.4-5**), secondary coma (horizontal and vertical, **Supplementary Fig.11-12**) and trefoil (oblique and vertical, **Supplementary Fig.7-8**) in biplane setup, as well as coma and secondary coma in astigmatism system. (2) Aberrations that act differently at various axial positions. The second category includes aberrations such as astigmatism (vertical and diagonal, **Fig.2a, Supplementary Fig.2-3**), secondary astigmatism (vertical and diagonal, **Supplementary Fig.9-10**), primary spherical (**Fig.2a, Supplementary Fig.6**), secondary spherical (**Supplementary Fig.13**) in biplane. Aside from these aberrations that are common to both systems, trefoil (oblique and vertical, **Supplementary Fig.7-8**) in the astigmatism system also falls into the second category. The result indicates that the correction of aberrations may not always benefit the attainable axial localization precision but rather depending on the specific type of aberration that is corrected. Specifically, the correction of aberrations from the first category is more likely to be beneficial for improving axial resolution across the axial imaging range, while the correction within the second category can either enhance or worsen the achievable resolution at different z positions, provided that an accurate *in situ* PSF is retrievable and is used during localization. For example, in biplane system, for coma aberration from the first category, as the aberration amplitude increases from 0 to 3 λ/2π, the axial localization precisions get uniformly deteriorated across the axial range, with the average precision value worsened from 440.0 nm to 797.8 nm (a 1.8x increase) (**Fig.2b**). However, instead of deteriorating the axial precision, astigmatism improves axial localization precision of a single molecule in a biplane setup within an axial range of −400 nm to 400 nm (**Fig.2a**), due to the extended focusing effect induced by astigmatism near focal plane. In addition, primary spherical aberration (**Fig.2a**) in general rapidly deteriorates axial precision but such deterioration is much more pronounced in negative direction of the focus, where the PSF remains Gaussian-shaped through an extended z-range comparing to the positive direction where PSF resembles concentric rings that rapidly evolve along the axial direction. Intuitively, comparing to the slowly changing Gaussian shapes, these rapidly evolving rings encode more information about the axial location of a single molecule, leading to better axial localization precision.

Among the investigated aberrations, primary spherical aberration exerts the most prominent axial resolution deterioration when measuring average achievable axial precision per photon count in the range of −600 to 600 nm (**Fig.2b, Supplementary Fig.15**) in both biplane and astigmatism systems. And coma is the next aberration in line with a significant effect on axial resolution. In contrast, increasing astigmatism has a negligible effect on average axial precision (**Fig. 2b**), noting that such effect is far from negligible when a particular axial location (instead of the mean) is examined (**Fig.2a**). In the case of primary spherical aberration, axial precision increases from 440 nm to 1352 nm (a 3x increase) in biplane, from 420 nm to 1380 nm (a 3.3x increase) in astigmatism setup as the aberration amplitude increases from 0 to 3 λ/2π (**Fig. 2b**). By contrast, the average axial precision remained nearly constant as astigmatism amplitude increases from 0 to 3 λ/2π.

**Fig. 2:**
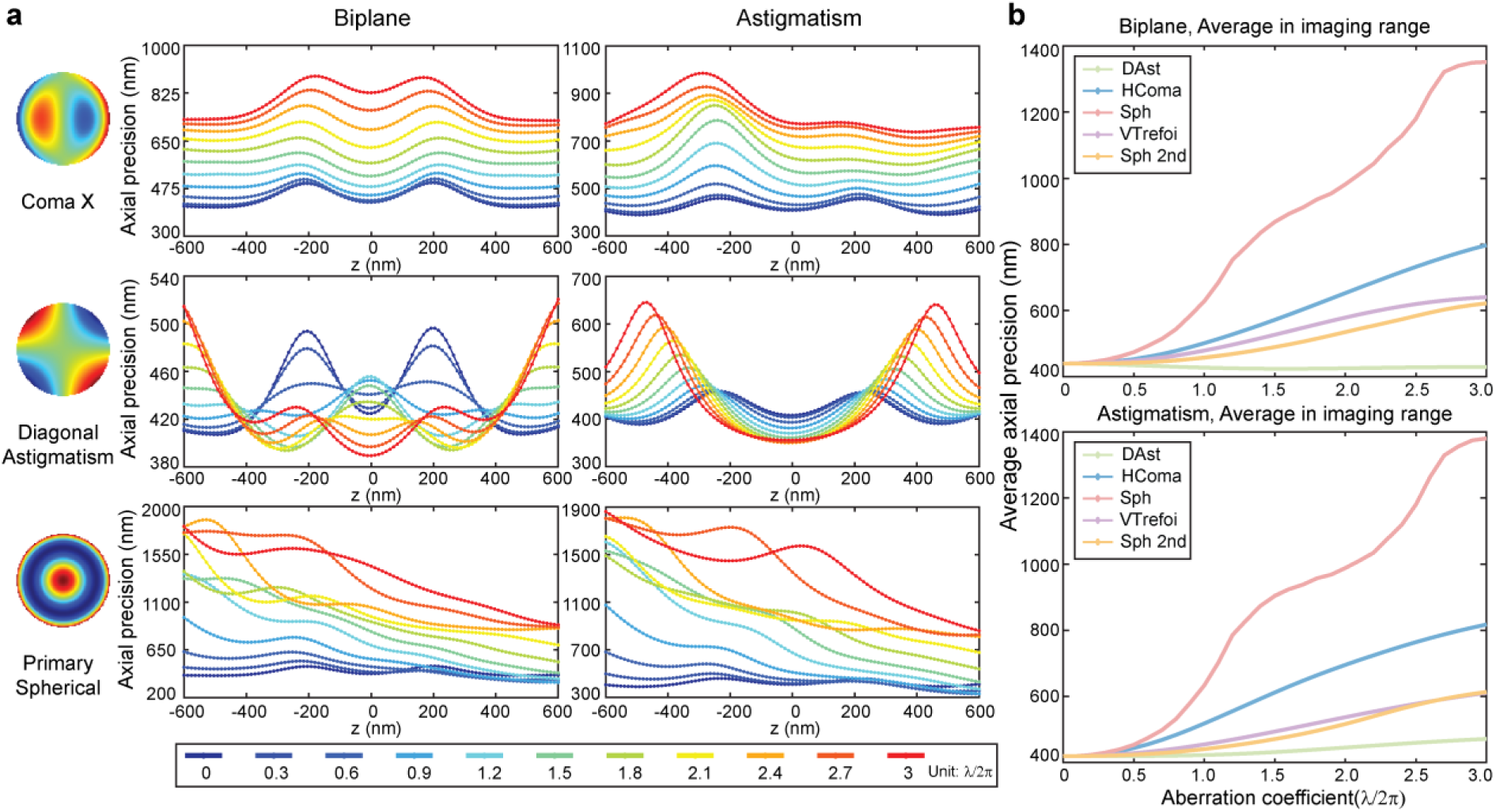
Aberration’s influences on axial localization precision. (a) Axial localization precision along the z-axis for horizontal coma, diagonal astigmatism, and spherical aberration in both biplane (left) and astigmatism (right) systems. Each figure is color-coded to represent aberration coefficients from 0 to 3 in increments of 0.3, measured in units of λ/2π. (b) Averaged axial localization precision with varying aberration types and amplitudes in biplane (top) and astigmatism (bottom) systems. The average is calculated from 101 frames within the −600 nm to 600 nm z-range with equal spacing. Abbreviations: DAst (diagonal astigmatism), HComa (horizontal coma), Sph (spherical), VTrefoil (vertical trefoil), Sph 2nd (secondary spherical). Simulation conditions remain consistent: *λ* = 680 nm, *I* = 0.5 photons/plane (biplane) and 1 photon (astigmatism), *bg* = 0, objective *NA* = 1.4, objective immersion medium refractive index (*n_obj_*) = 1.52, biplane distance: 400 nm, astigmatism amplitude: 1.4 λ/2π.

### Localization precision in presence of refractive index mismatch

Oil immersion objectives are commonly used in single molecule imaging due to their high numerical aperture. While the samples are usually mounted in water-based medium, the refractive index mismatch between the sample medium and objective immersion medium results in a specific type of aberration, commonly referred as index mismatch (IMM) aberration^32^ (**Fig. 3a**). IMM aberration induces focal shift, causing a depth-dependent displacement between intended and actual focal positions and has been experimentally characterized^41^. IMM aberration also distorts the PSF shape, reduces the image contrast, resulting in degraded localization precision both laterally and axially^18^. The extent of the focal shift and degradation depends on both the differences in refractive indices and imaging depth. In this section, we assess the average achievable localization precision around the actual focal position under varying refractive indices of the sample medium and at different depths.

**Fig. 3:**
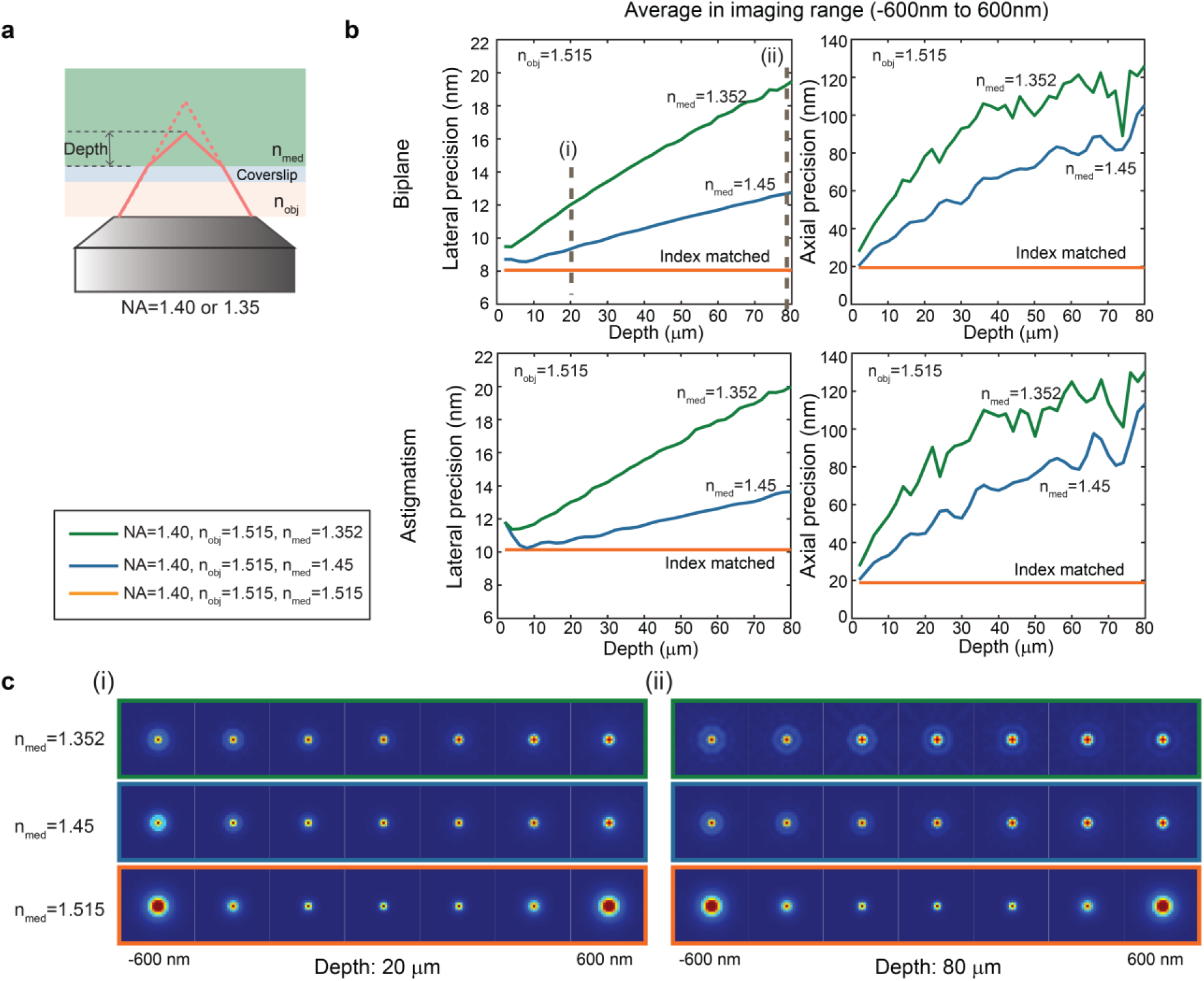
Single molecule localization precision in index mismatch (IMM) scenarios. (a) Schematic illustrating refractive index mismatch. (b) Lateral (left) and axial (right) localization precision at different imaging depths under index-mismatched conditions in the biplane (top) and astigmatism (bottom) systems. NA: numerical aperture, n_obj_: refractive index of the objective immersion medium, n_med_: refractive index of the sample immersion medium. (c) Example Point Spread Functions (PSFs) within the −600 nm to 600 nm range corresponding to (i) depth = 20 μm and (ii) depth = 80 μm cases, as indicated in figure (b). Simulation conditions: *λ* = 680 nm, *I* = 500 photons/plane for biplane and *I* = 1000 photons for astigmatism system, *bg* = 10 photons, objective *NA* = 1.4, objective immersion medium refractive index (*n_obj_*) = 1.515, biplane distance: 400 nm, astigmatism amplitude: 1.4 λ/2π. Imaging depth ranges from 2 μm to 80 μm with a 2 μm step size. At each depth, 101 frames were generated over the z-range from –600 to 600 nm relative to the actual focal position, with equal spacing, and the average values were calculated. The region of interest (ROI) size is 32 × 32 pixels.

IMM affected PSF can be modeled by adding a depth dependent and rotationally symmetric pupil phase derived from the Gibson-Lanni’s model in the pupil function^33^. Assuming that the refractive indices of the coverslip and the objective immersion medium are the same, such IMM phase can be expressed analytically as^20^:

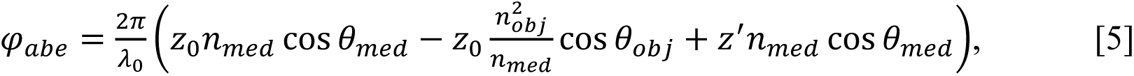

where 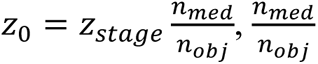 is a correction factor which calucated by the ratio of refractive index of sample medium *n*_*med*_ to the refractive index of objective immersion medium *n*_*obj*_, *z*_*stage*_ is the piezo stage position with its zero value attained when focusing the objective on the coverslip surface, *z*^′^ is the relative defocus position of a single molecule with respect to *z*_0_, *λ*_0_ is the wavelength of fluorescence emission in vacuum. Considering the focal shift casued by this IMM induced aberration which is similar to spherical aberration, we need to shift the imaging range accordingly. Similar to our approach for spehrical aberration, we selected the axial imaging range where the PSFs largely retained their focused characteristics based on the peak intensity of the PSFs to evaluate the localization precision (**Methods**).

We examined cases using a high-NA oil immerion objective with 1.4 NA in combination with various sample mounting media with different refractive indices, including water-based dSTORM buffer (index of refraction: 1.352), glycerol-based Vectashield^42^ (index of refraction: 1.45) and oil index-matched buffer^43^ (index of refraction: 1.515). The index mismatch exacerbates localization precisions both laterally and axially, and such effect is much more significant axially than laterally in both biplane and astigmatism setup (**Fig. 3b**). For instance, in biplane setup, when imaging single molecules in dSTORM buffer with a refractive index of 1.352 and using an oil immersion objective with a refractive index of 1.515 at depths ranging from 0 µm to 80 µm, the lateral localization precision rises from 9.5 nm to 19.5 nm, representing a 2.1x increase, while the axial localization precision gets deteriorated from 27.8 nm to 126.1 nm, showing a 4.5x increase. This is because in addition to blurring the emission pattern of single molecule which degrades both lateral and axial precision, IMM reduces the PSF shape modulation amplitude along the axial direction, as demonstrated by the PSFs in **Fig.3c**, significantly reducing the Fisher information content along the axial direction. Such demodulation effect, however, does not influence the achievable lateral precision.

This resolution worsening effect can be alleviated by minimizing the refractive index difference between objective immersion medium and sample’s mounting medium. Various approaches have been introduced to increase the medium’s refractive index, including the addition of 2,2-thiodiethanol(TDE)^44^, glycerol^45^, and more recently, 3-pyridinemethanol (3-PM)^43^, which can match the oil index of 1.515. In the case of using Vectashield with higher index of refraction 1.45, at a depth of 80 µm in biplane setup, the localization precision was 12.7 nm laterally and 105.3 nm axially, showing a 1.5x and 1.2x improvement compared to dSTORM buffer (**Fig.3b**). When the refractive index is ideally matched, precision should not degrade when imaging deep into the specimen as shown in our simulations, assuming no contributions from aberrations caused by the imaging system or refractive index inhomogeneity within the biological sample. While objectives are typically chosen based on designed theoretical achievable resolution, aligning the refractive index of the sample mounting medium with the objective’s immersion medium serves as an important strategy to mitigate refractive index mismatches aberrations and substantially enhance localization precision.

## WAVEFRONT ESTIMATION IN PRESENCE OF EXISTING ABERRATIONS

Most system induced and sample induced aberrations, characterized by aberrated wavefront in the pupil plane, can be corrected using adaptive optics (AO)^16^ by measuring and subsequently compensating the concurrent distortion. To estimate these wavefront distortions, single molecule emission patterns which are routinely recorded during SMLM experiments represents an unique and convenient source as their shape contains abundant information about existing aberrations in both the imaging system and the specimen^13,26^. For each single molecule emission pattern, the wavefront estimation precision depends on its signal to background ratio, axial position as well as the complex shape formed by different aberration types and amplitudes. In this section, we explored the achievable precision limits for wavefront estimation based on single molecule emission patterns in a 3D imaging system.

### Wavefront estimation precision along the axial position

Common among all aberration-modes investigated, the achievable wavefront measurement precision from single emitters varies significantly at different axial positions. Methodologies have been developed to reference the emission patterns of single molecules to acquire wavefront information^13,26^. This leads to the question that the PSFs from which axial positions contain a greater amount of information about the wavefront, which in turn could potentially result in more accurate estimations of the wavefront shape. In the presence of a small amount of aberration, the measurement of aberrations using out-of-focus emission patterns generally yields better precision than using in-focus ones (**Supplementary Fig.16-17**). This can be intuitively understood because, when aberrations are minor, the out-of-focus PSFs may exhibit more distinct characteristics, aiding in the estimation of the aberrations. This behavior, however, is reversed as wavefront distortions increase (**Fig.4, Supplementary Fig.16-17**). At an aberration amplitude of 3 λ/2π, two categories of aberrations exhibit different trends at different axial positions. The first group includes astigmatism, coma, trefoil, secondary astigmatism, and secondary coma, for which in-focus PSFs yield better aberration estimation precision. For example, when measuring coma with an amplitude of 3 λ/2π in biplane system, the measurement uncertainty is 1.4 times better for in-focus PSFs compared to PSFs situated 600 nm away from the focus. Similarly, when measuring trefoil in astigmatism system at an amplitude of 3 λ/2π, the precision for in-focus PSFs is also 1.3 times better than out-of-focus PSFs. The secondary category comprises primary spherical and secondary spherical aberrations. With substantial spherical distortion, a scenario commonly encountered when imaging through whole cells and tissues or in cleared/expanded specimens with high-NA objectives, the measurement precision of spherical aberrations is more favorable on the positive side of the PSF, where concentric ring patterns are formed, rather than in the negative direction, which consists of gradually changing Gaussian-shaped PSFs. This is in concert with the previously discussed behavior of axial localization precision in the presence of spherical aberration. In both cases, estimation precisions benefit from the rapidly evolving ring structure on the positive side generating increased amount of Fisher information than its Gaussian counterpart despite its considerably larger emission pattern. In biplane setup, the estimation precision of large spherical at +600 nm is 1.6 times better than at −600 nm and the estimation for secondary spherical aberration is 1.9 times better in the same context.

**Fig. 4:**
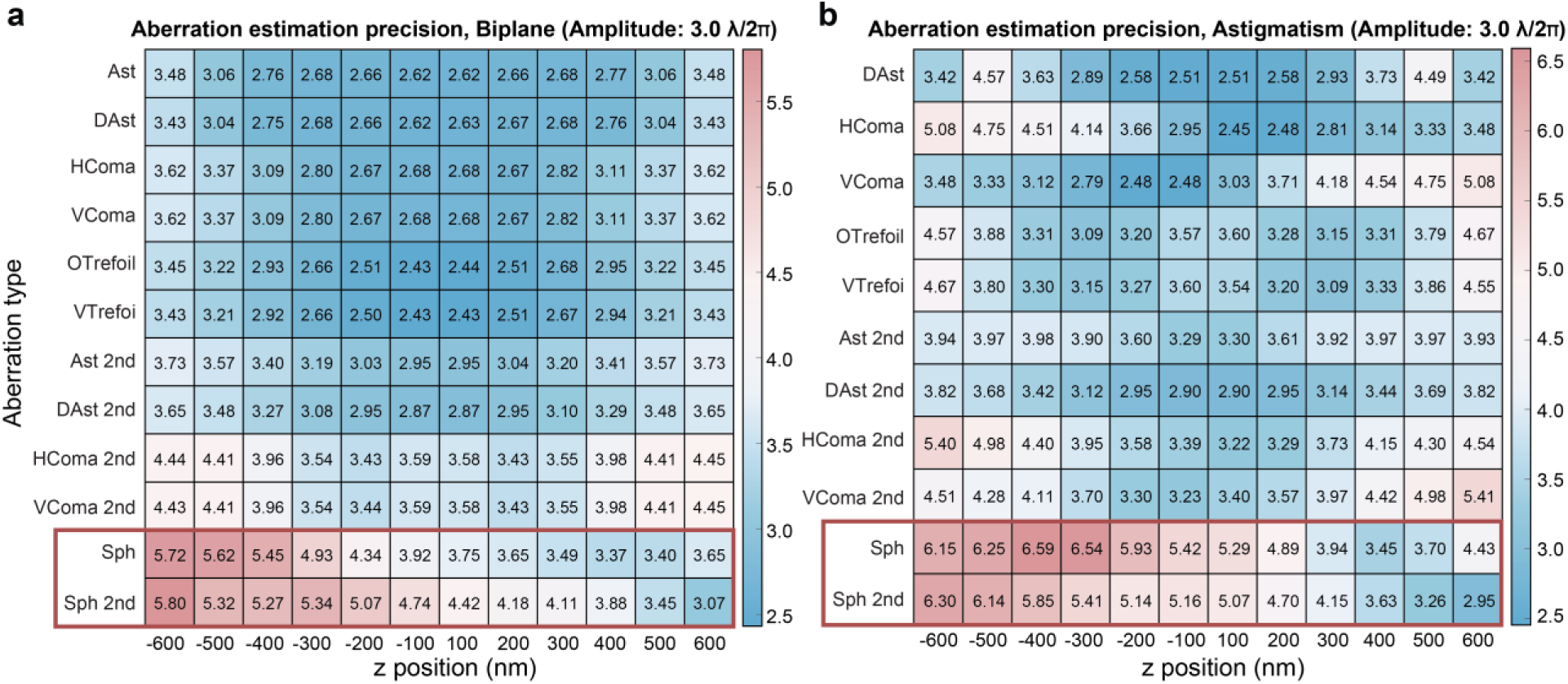
Aberration estimation precision for various aberration modes at different axial positions. Aberration estimation precision in the presence of 3 λ/2π aberration in the (a) biplane and (b) astigmatism system. The values in each cell represent the average precision calculated within a 100 nm range. Abbreviations: Ast (astigmatism), DAst (diagonal astigmatism), HComa (horizontal coma), VComa (vertical coma), Sph (spherical), OTrefoil (oblique trefoil), VTrefoil (vertical trefoil), Ast 2nd (secondary astigmatism), DAst 2nd (secondary diagonal astigmatism), HComa 2nd (secondary horizontal coma), VComa 2nd (secondary vertical coma), Sph 2nd (secondary spherical). Simulation conditions: *λ* = 680 nm, *I* = 0.5 photons/plane for the biplane system, *I* = 1 photon for the astigmatism system, *bg* = 0, *NA* = 1.4, *n_obj_* = 1.52, biplane distance: 400 nm, astigmatism amplitude: 1.4 λ/2π.

### Estimation crosstalk among different aberration types

Wavefront distortions generated by the imaging system and the specimen are often complex, consisting of multiple types of aberration modes when described using Zernike polynomials. These distorted wavefronts manifest aberrated PSFs with specific features carrying information of the wavefront shape. The features characterizing one aberration mode can be weakened or sometimes strengthened in presence of another aberration, which in turn deteriorates or improves the wavefront measurement precision. Here, we investigated the measurement interaction between aberration modes, specifically examining how the presence of one aberration mode influences the estimation precision of another.

To provide a summary of the collective impact on aberration estimation precision, we computed the average precision across an axial range from −600 nm to 600 nm. By altering the amplitudes of pre-existing aberration modes, we calculated the attainable precisions in measuring a target aberration mode at a specific base amplitude. In this case, we illustrate this with an example of a base amplitude set at 2 λ/2π. Generally, most aberrations tend to reduce the precision of estimating other aberrations in both biplane and astigmatism systems, although at varying rates (see **Fig. 5, Supplementary Fig.18-23**). Some aberrations, on the other hand, have minimal effects or can even enhance the precision of estimating other aberrations. For example, in both biplane and astigmatism system, increasing coma degrades the precision of astigmatism estimation but has minimal impact on estimating spherical aberration (**Fig. 5, Supplementary Fig.19**). This can be intuitively explained by observing the aberrated PSFs (**Fig. 5a** (i) to (iv)): Significant coma conceals astigmatism features, but it does not obscure the large spherical features. When a large coma is present, spherical rings are still visible. Diagonal astigmatism presence enhances the precision of spherical and secondary coma aberration estimation in the biplane system (**Fig. 5, Supplementary Fig.18a)**. In the astigmatism system, diagonal astigmatism improves the estimation precision for most aberrations except secondary diagonal astigmatism (**Supplementary Fig.18b**). For instance, in the biplane system, introducing diagonal astigmatism results in an 18% improvement in spherical aberration estimation precision. In the astigmatism system, trefoil estimation precision improves by 14%, and spherical aberration estimation by 12%. These observations indicate that the existence of astigmatism either intentionally or not actually helps to encode an increased amount of information in the emission patterns and thus benefit wavefront estimation from single molecules. Among the tested aberration modes, spherical aberration has the most pronounced effect on aberration measurement (**Fig. 5, Supplementary Fig.20**). For instance, an increase in spherical aberration of 3 λ/2π from a baseline of 0 results in a 4.7-fold worsening of achievable precision for diagonal astigmatism in the biplane system and a 4.3-fold worsening in the astigmatism system. This phenomenon is likely attributed to the axial spread of the spherically-aberrated PSFs, which obscures the characteristic shapes formed by other aberrations (**Fig. 5a,b** (vii), (viii)). Similar observations can be made for secondary spherical aberration (**Supplementary Fig.20**). Given the limited signal-to-background ratio in single-molecule datasets, these findings underscore the importance of prioritizing the correction or mitigation of spherical aberrations to restore the information content of other aberration modes carried by the emission pattern.

**Fig. 5:**
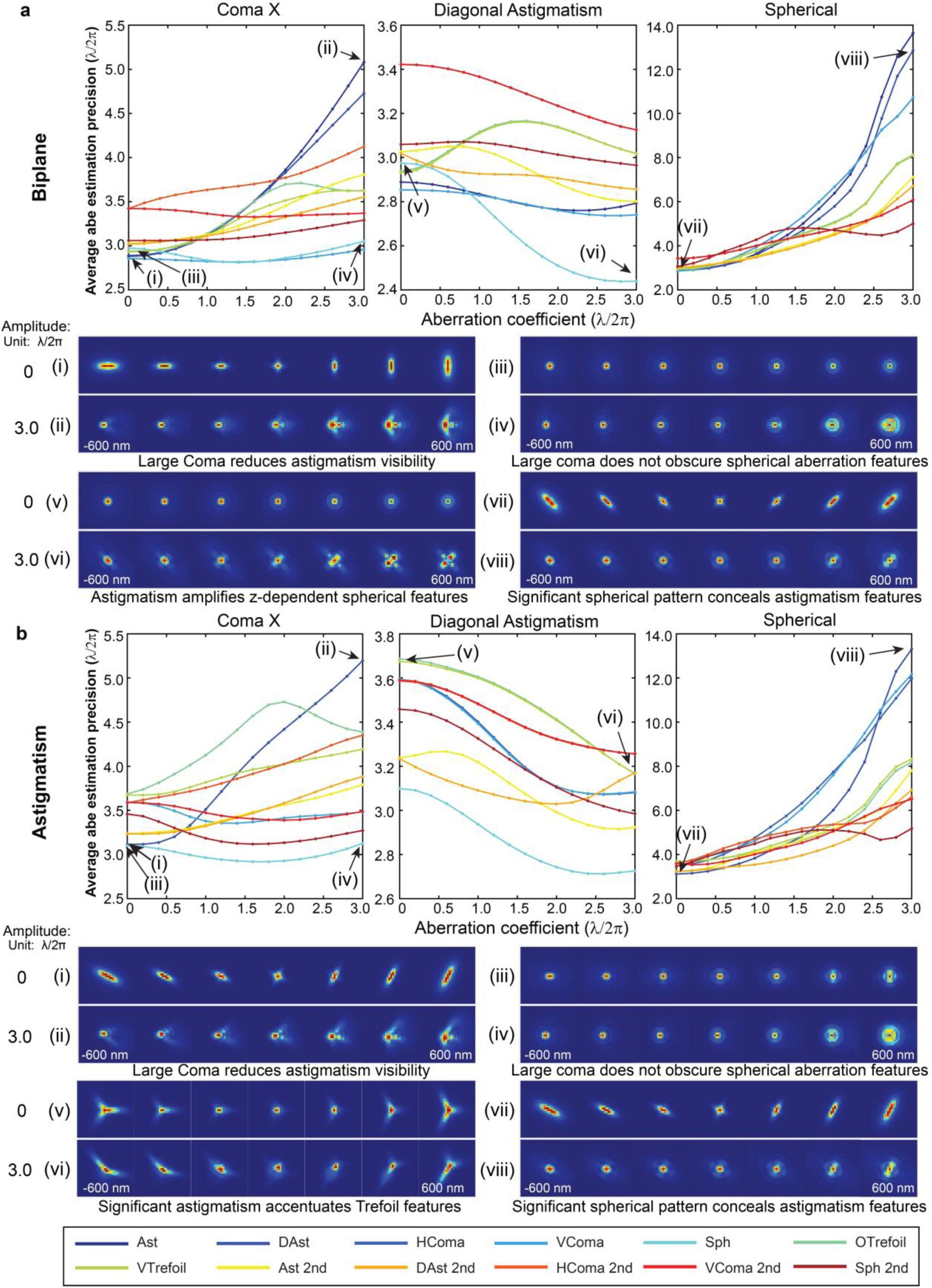
Aberration estimation precision in the presence of preexisting aberrations. (a) Precision of aberration estimation when estimating each aberration at an amplitude of 2 λ/2π while introducing varying levels of another aberration (horizontal coma, diagonal astigmatism, and spherical aberration) in biplane system. The precision values are calculated by averaging within the −600 nm to 600 nm z range. The rainbow colors represent different aberration modes, ranging from vertical astigmatism to secondary spherical aberration (Wyant order). (i) to (viii): Examples of PSFs within the −600 nm to 600 nm range, corresponding to cases as indicated in the figures. (b) The same analysis as in (a) but focusing on the astigmatism system. (i) to (viii): Examples of PSFs within the −600 nm to 600 nm range, corresponding to cases as indicated in the figures. Simulation conditions: *λ* = 680 nm, *I* = 0.5 photons/plane for the biplane system, *I* = 1 photon for the astigmatism system, *bg* = 0, objective *NA* = 1.4, *n_obj_* = 1.52, biplane distance: 400 nm, astigmatism amplitude: 1.4 λ/2π.

## DISCUSSION

In this study, we conducted a systematic analysis of the achievable estimation lower bounds for localization precision per photon count in both the lateral and axial directions, as well as the precision of aberration estimation in the presence of aberrated single molecule emission patterns in both biplane and astigmatism setups. Our findings shed light on the intricate relationship between aberrations and the performance of 3D single molecule localization techniques. Our observations indicated that, in general, aberrations tend to have mixed effects on localization precision. While they often deteriorate lateral precision near the focus, it is important to note that some aberrations can surprisingly enhance the precision of out-of-focus molecules. This duality suggests a complex interplay between aberrations and the characteristics of single molecule emission patterns. Additionally, our investigation revealed the significant alteration in the uniformity of achievable lateral localization precision across an axial range, which varies with the degree of aberrations present. This observation holds particular significance for researchers engaged in imaging through thick samples, where maintaining uniform precision across the entire depth range is crucial. In examining axial localization precision, we found that primary spherical aberration exerted a dominant influence, significantly deteriorating axial precision. On the other hand, the presence of astigmatism was associated with improved axial precision, especially within a typical range close to the focal plane. These observations underscore the necessity of considering the specific aberrations at play to optimize axial precision in SMLM. We also observed interactions between different aberrations. Some aberrations may enhance or hinder the estimation precision of other aberrations. Notably, pre-existing astigmatism appeared to facilitate the estimation of spherical aberration, whereas pre-existing spherical aberration had a significant impact on the estimation of other aberrations. This insight highlights the need for careful consideration of aberration combinations and their potential influence on estimation accuracy.

Noting that these bounds were calculated in the scope of ideal case with no background, no noise and single photon count, in practice, a new set of precision limits could be obtained from different settings of modality parameters: intensity, background, wavelength, NA, etc. Our Fisher information-based analyses here offer a theoretical framework for understanding the behavior of different aberration types in SMLM. These findings can serve as a valuable reference for aberration correction strategies, prioritizing the correction of the most detrimental aberrations, and exploring the introduction of beneficial aberrations to enhance estimation. This analysis holds the potential to inform the design of optical systems, the optimization of imaging conditions, and the development of effective aberration control technologies in the realm of ultrahigh-resolution optical imaging based on single molecule detections.

## Acknowledgments

We would like to thank Sheng Liu, Peiyi Zhang, Donghan Ma, Fan Xu for the discussions on the topic. We thank Peiyi Zhang and Sheng Liu for suggestions on PSF simulations. This work was supported by the National Institute of General Medical Sciences (R35GM119785).

## Data and code availability

The data and code that support the findings of this study are available from the authors upon request.

## Methods

### PSF generation

We generated the PSFs based on scalar diffraction theory. We calculated the Point spread function (PSF) of an imaging system from the Fourier transform of the pupil function^46^:

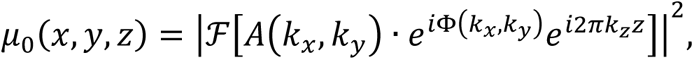

where *μ*_0_(*x*, *y*, *z*) represents the PSF intensity at position (*x*, *y*, *z*) in image plane, ℱ denotes the Fourier transform, *A*(*k*_*x*_, *k*_*y*_) · *e*^*i*Φ(*k*_*x*_,*k*_*y*_)^ is the pupil function, *A*(*k*_*x*_, *k*_*y*_) is the magnitude and Φ(*k*_*x*_, *k*_*y*_) is the phase of the pupil function, *e*^*i*2*πk*_*z*_*z*^ describe the defocus phase, with *k*_*z*_ = 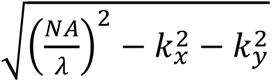, NA is the numerical aperture of the objective lens, and *λ* is the emission wavelength.

Aberrations were modeled as the phase deviation in the pupil plane, represented as a combination of a series of Zernike polynomials^47^:

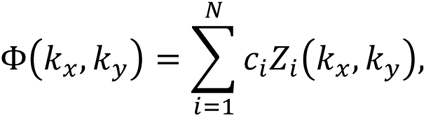

where *Z* _*i*_(*k*_*x*_, *k*_*y*_) is the *i*th Zernike mode, *c*_*i*_is the corresponding coefficient, *N* is the number of Zernike modes we considered. In our simulation, we considered Zernike aberrations from vertical astigmatism to secondary spherical (N=5 to N=16, Wyant order). For index mismatch situation, we add an additional aberration phase to the pupil function as described in equation [5].

We utilized a custom PSF generation toolbox in MATLAB to generate PSFs^13^. In the case of the astigmatism system, a fixed vertical astigmatism of 1.4 λ/2π (*c*_5_ = 1.4) was introduced into the pupil function (**Supplementary Fig. 1**). The pupil function was then modified based on the simulation conditions, and the normalized PSF was generated. Subsequently, the PSFs were adjusted for intensity and background using the formula: *μ* = *I* ∗ *μ*_0_ + *bg*, where *μ*_0_ is the normalized PSF, *I* represents the intensity, and *bg* signified the background. In the case of the biplane setup, we configured the Zernike coefficients for the pupil function and generated a pair of PSFs based on the constructed pupil function at specified positions (*x*, *y*, *z*_1_) and (*x*, *y*, *z*_2_), with a predefined biplane distance of *z*_2_ − *z*_1_ = 400 *nm* (**Supplementary Fig. 1**).

### Calculation of Fisher information and Cramer-Rao Lower Bound in SMLM

In the case of single molecule emission pattern detection, the probability of photon detection in each pixel follows Poisson distribution. Under Poisson distribution, the numerical expression of Fisher information in SMLM can be simplified as follows^20^:

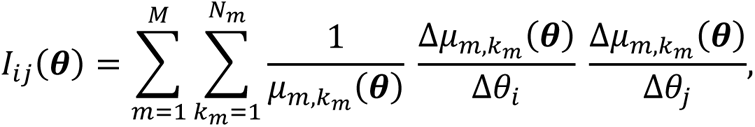

Where *M* is the number of planes, equal to 1 for astigmatism system, 2 for biplane system; *N_m_* is the pixel numbers in the *m*th plane, in our simulation we used 32 × 32 pixels subregions, so *N_m_* = 1024; *μ*_*m*,*km*_(***θ***) is the value of the PSF model at pixel *k* in *m*th plane; ***θ*** is the parameter vector to be estimated. In the context of assessing the impact of aberrations on localization precision, the parameters of interest include {*x*, *y*, *z*, *I*, *bg*}, where {*x*, *y*, *z*} is the position of the single molecule, *I* is intensity, *bg* is the background noise. When calculating the derivative numerically, we set the increments Δ*x* = 0.1 *pixel*, Δ*y* = 0.1 *pixel*, Δ*z* = 0.01 *μm*. For evaluating the precision of aberration estimation, we considered Zernike modes from astigmatism to secondary spherical, the parameter vector in this case includes {*x*, *y*, *z*, *I*, *b*, *c*_5_, *c*_6_ … *c*_*i*_, … *c*_16_}, with each aberration coefficient changing by Δ*c* = 0.01 when calculating derivatives. Consequently, the Fisher information matrix was a 17 ×17 matrix. The diagonal terms of the inverse of Fisher information matrix correspond to the CRLB for the respective parameter. The square root of the diagonal terms in the CRLB matrix provided the theoretical achievable estimation precision for each parameter.

### Focal plane determination

In cases where the imaging system experiences spherical aberration or index mismatch aberration, it induces an axial displacement of Point Spread Functions (PSFs). This shift results in a deviation of the actual focal plane from its ideal position. In practical applications, we typically acquire PSFs within the range where they exhibit their highest peak intensity. So, we defined the actual focal plane as the plane around which PSFs display their maximum peak intensity. Specifically, within a range of [-600, 600] nm relative to the defined actual focal plane, the PSFs consistently exhibit peak intensity.

To determine this actual focal plane, we systematically generated PSFs over a range of −3 to 3 µm relative to the ideal focal position, with a precise step size of 10 nm. Subsequently, we constructed a curve representing the peak intensity of the PSFs along the z-direction. We then calculated the sum of peak intensity values within a 1200 nm window that slide along the range from −3 µm to 3 µm. The central position of this window, where the sum of peak intensity reached its maximum value, was identified as the actual focal position. This window, along with its associated range, formed our simulated axial range. Further details regarding the focal shift due to spherical aberration and index mismatch aberration are presented in **Supplementary Fig. 24**.

### Data simulation

We used customized PSF toolbox in MATLAB to tune the coefficients of Zernike polynomials to modify the phase aberration. We then generated the PSFs and calculated the CRLB and estimation precisions based on the methods discussed above. In all our simulations, the wavelength remained at 680 nm, image size was 32 × 32 pixels with a pixel size of 130 nm. The biplane distance was set at 400 nm, while the astigmatism amount for astigmatism system was 1.4 λ/2π. The investigated axial range was from −600 nm to 600 nm relative to the defined actual focal position. We studied 12 typical aberration modes (from astigmatism to secondary spherical, Wyant order) When investigating the effects of aberration modes on localization precision (**Fig.1-2),** we simulated the situation of imaging with oil-immersion, omitting index mismatch, and with normalized intensity. We set the NA of objective to be 1.4, the refractive index of the objective immersion medium to be 1.52, intensity to be 0.5 photons/plane for biplane and 1 photon for astigmatism and background noise to be 0. We studied 12 typical aberration modes (from astigmatism to secondary spherical, Wyant order) with each coefficient ranges from 0 to 3 λ/2π, with step size 0.3 λ/2π. For each aberration situation, we simulated 101 frames over the axial range from −600 nm to 600 nm with respect to the defined actual focal position with equal spacing distance and calculated the square root of CRLB to represent the localization precision at each position. Lateral localization precision was defined as the root mean square (RMS) of x- and y-localization precision, whereas axial localization precision focused on z-localization precision.

When simulating index mismatch situation (**Fig.3**), we considered the use of oil immersion objectives (*NA = 1.40, n*_*obj*_ =*1.515*) with dSTORM buffer (*n*_*med*_ =*1.352*) and glycerol-based Vectashield (*n*_*med*_ =*1.45*) and oil index-matched buffer (*n*_*med*_ =*1.515*). The intensity was set at 500 photons/plane for the biplane system, with background noise at 5 photons/pixel per plane, and for the astigmatism system, intensity was 1000 photons with background at 10 photons/pixel. In each situation, we modified the pupil function to account for the corresponding index mismatch aberration at different depths, from 0 μm to 80 μm with step size 2 μm. At each depth, we simulated 101 frames over the axial range from −600 nm to 600 nm relative to the defined actual focal position with equal spacing distance and calculated the lateral and axial localization precision at each position. Then we took the average value for lateral and axial localization precision respectively in the imaging range as the localization precision at each depth.

When studying the precision of aberration estimation along the z-direction, we maintained the same simulation conditions used to investigate the effects of aberration on localization precisions, but introduced additional estimating parameters: {*x*, *y*, *z*, *I*, *b*, *c*_5_, *c*_6_ … *c*_*i*_, … *c*_16_} into the Fisher information and CLRB matrices. We simulated 101 frames over the axial range from −600 nm to 600 nm with respect to the defined focal position with equal spacing distance and calculated the square root of CRLB to represent the estimation precision at each position. The values in **Fig.4** were calculated as the average estimation precision for every 100 nm.

When investigating the crosstalk between aberrations, we set the coefficient of the target aberration mode to be 2 λ/2π and introduced various preexisting modes with different coefficients, ranging from 0 to 3 λ/2π with step size 0.2 λ/2π. For each coefficient, we simulated 101 frames across the axial range from −600 nm to 600 nm relative to the focal position with equal spacing and calculated the average estimation precision of the target aberration mode and compared the estimation precision with differing amounts of other aberrations.

