## Supplementary Materials for "Measurement precision bounds on aberrated single molecule emission patterns"

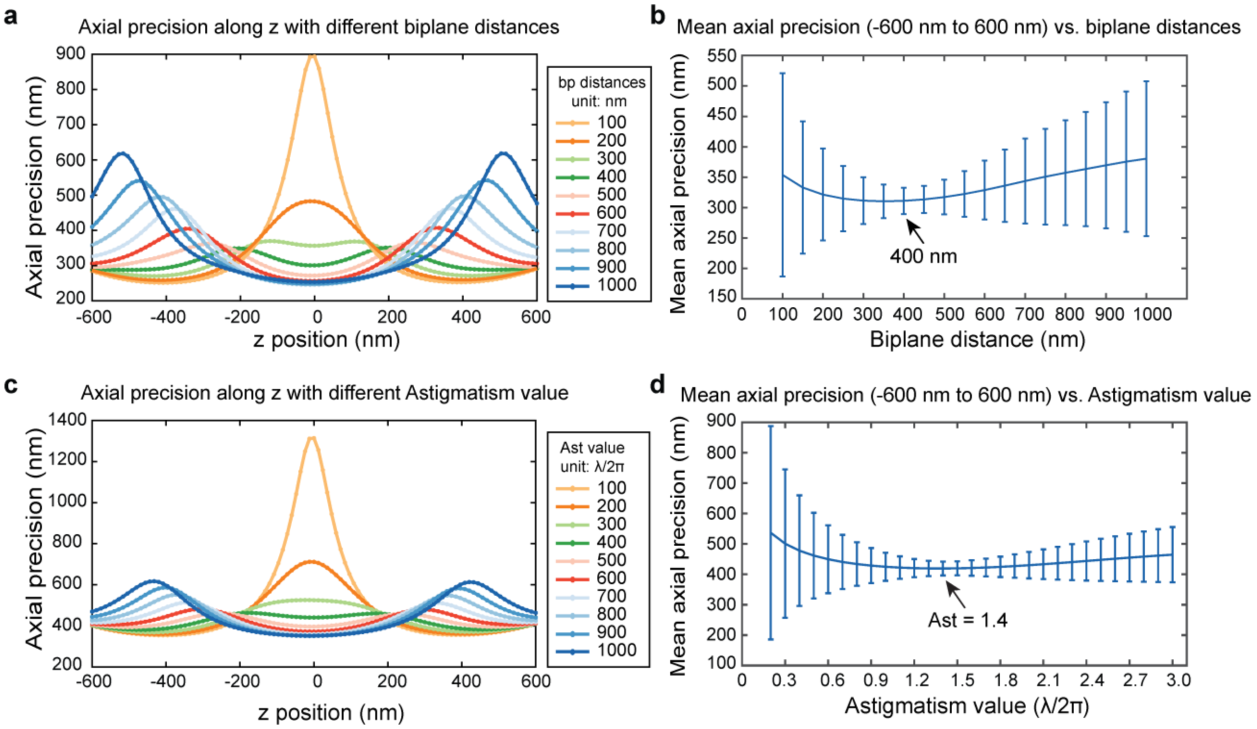

**Supplementary Fig. 1: Axial localization precision with different biplane distances and astigmatism values**. (a) Axial localization precision along the z-axis with varying biplane distances. (b) Mean axial localization precision with error bars within -600 nm to 600 nm z range vs. biplane distances. (c) Axial localization precision along the z-axis with varying astigmatism values. (d) Mean axial localization precision with error bars within -600 nm to 600 nm z range vs. astigmatism values. Simulation conditions: Wavelength ($\lambda$) = 680 nm, intensity ($I$) = 1 photon, background ($bg$) = 0, Numerical Aperture ($NA$) = 1.4, immersion medium with refractive index ($n_{obj}$) = 1.52, pixel size = 130 nm. The region of interest (ROI) size was set to 32 × 32 pixels. The optimal biplane distance and astigmatism value for maintaining uniformly high axial localization precision along the z-axis were chosen as biplane distance = 400 nm and astigmatism value = 1.4 λ/2π.

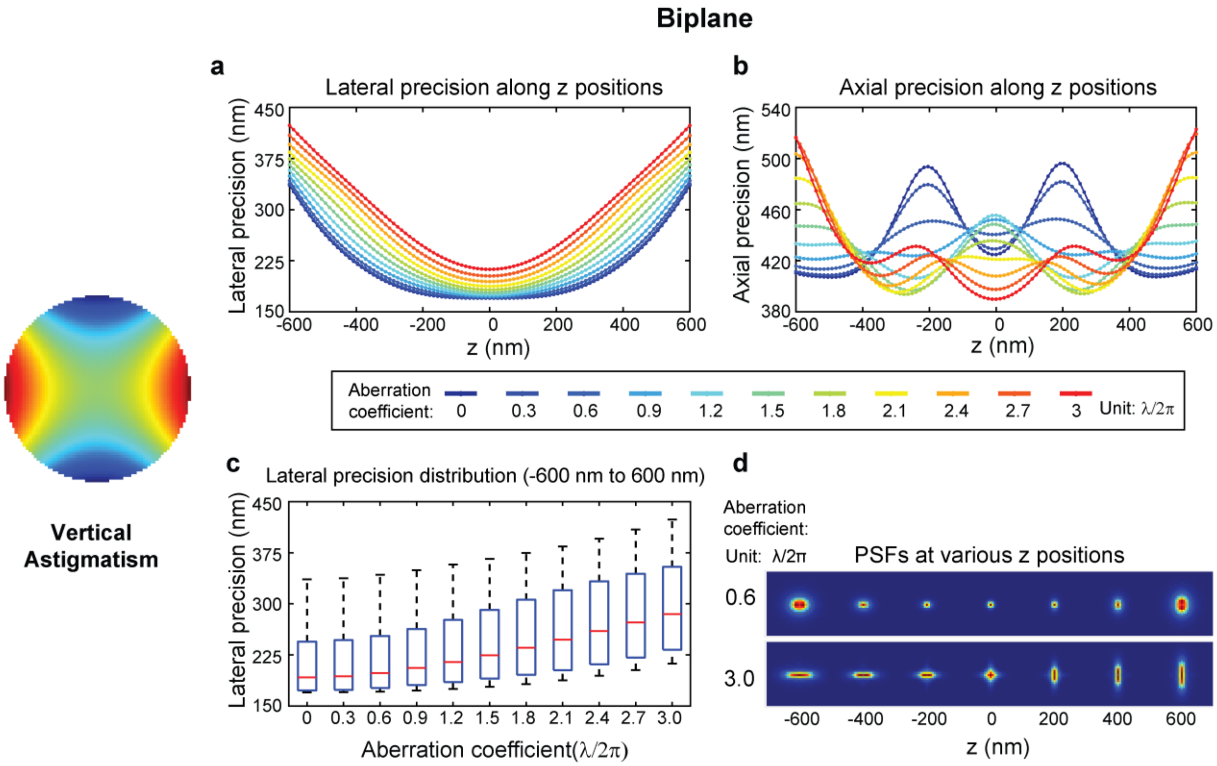

**Supplementary Fig. 2: Effects of vertical astigmatism on localization precision with varying amplitudes in biplane systems**. (a) Lateral localization precision along the z-axis with varying amplitudes of vertical astigmatism. The precision values are color-coded to represent aberration coefficients ranging from 0 to 3, with a step size of 0.3, measured in units of λ/2π. (b) Axial localization precision along the z-axis, similarly color-coded with varying vertical astigmatism amplitudes. (c) Distributions of lateral localization precision within the z range of -600 nm to 600 nm with varying amplitudes (ranging from 0 to 3). Each boxplot displays the median, 25th, and 75th percentiles of the data, while whiskers extend to non-outlier extreme points. Outliers are individually marked with plus signs. (d) Examples of PSFs at various z positions with small aberration (0.6 λ/2π) and large aberration (3.0 λ/2π). Simulation conditions: $\lambda$ = 680 nm, $I$ = 0.5 photons/plane for the biplane system, $bg$ = 0, $NA$ = 1.4, $n_{obj}$ = 1.52, pixel size = 130 nm, biplane distance = 400 nm. Lateral localization is calculated as the root mean square (RMS) of x- and y- localization precision. The region of interest (ROI) size was set to 32 × 32 pixels.

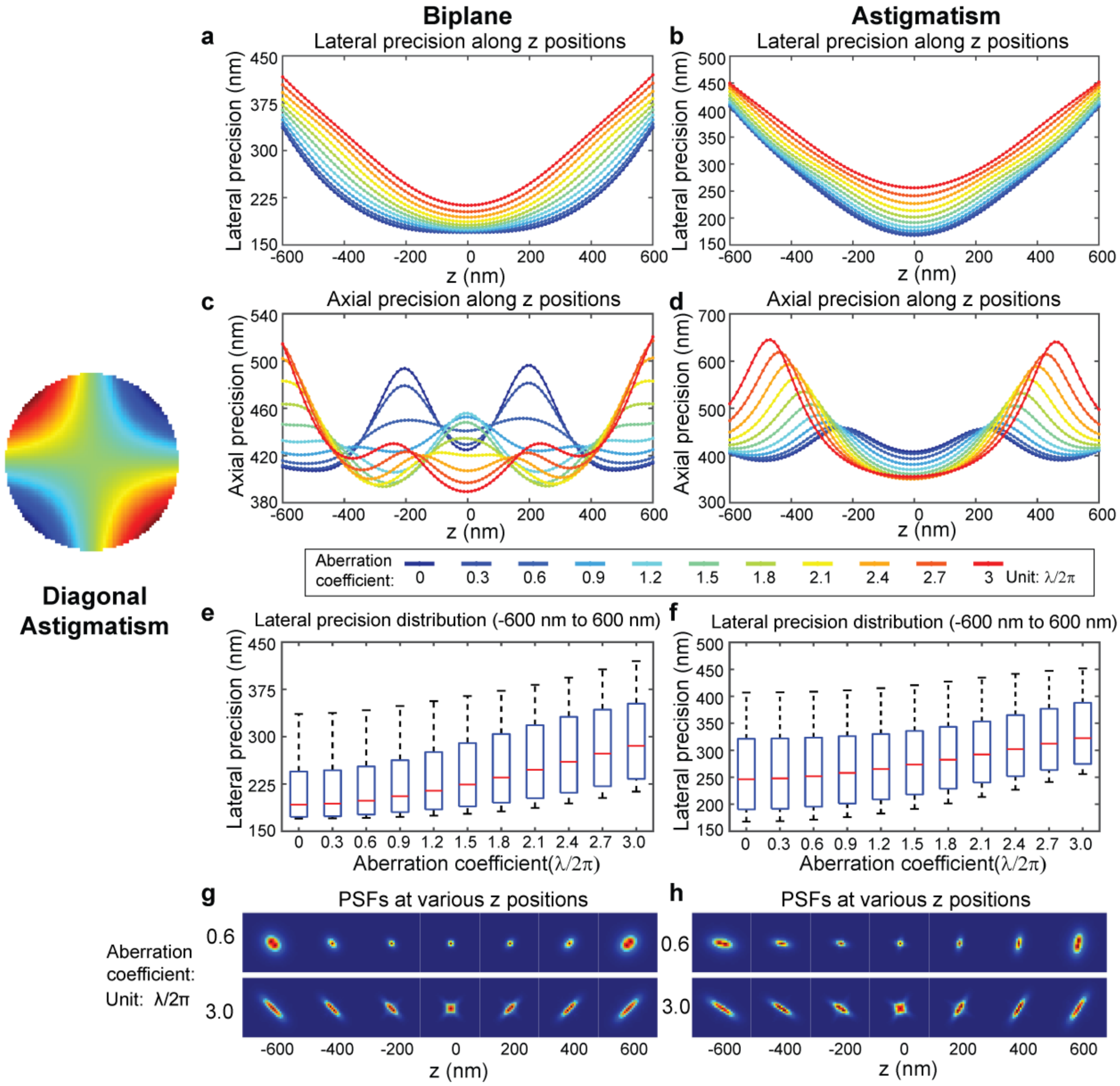

**Supplementary Fig. 3: Effects of diagonal astigmatism on localization precision with varying amplitudes**. (a, b) Lateral localization precision along the z-axis with varying amplitudes of diagonal astigmatism in biplane and astigmatism systems. The precision values are color-coded to represent aberration coefficients ranging from 0 to 3, with a step size of 0.3, measured in units of λ/2π. (c, d) Axial localization precision along the z-axis, similarly color-coded with varying diagonal astigmatism amplitudes in biplane and astigmatism systems. (e, f) Distributions of lateral localization precision within the z range of -600 nm to 600 nm with varying amplitudes (ranging from 0 to 3) in biplane and astigmatism systems. Each boxplot displays the median, 25th, and 75th percentiles of the data, while whiskers extend to non-outlier extreme points. Outliers are individually marked with plus signs. Simulation conditions: $\lambda$ = 680 nm, $I$ = 0.5 photons/plane for the biplane system, $I$ = 1 photon for the astigmatism system, $bg$ = 0, $NA$ = 1.4, $n_{obj}$ = 1.52, pixel size = 130 nm, biplane distance = 400 nm, astigmatism value = 1.4 λ/2π. Lateral localization is calculated as the root mean square (RMS) of x- and y- localization precision. The region of interest (ROI) size was set to 32 × 32 pixels.

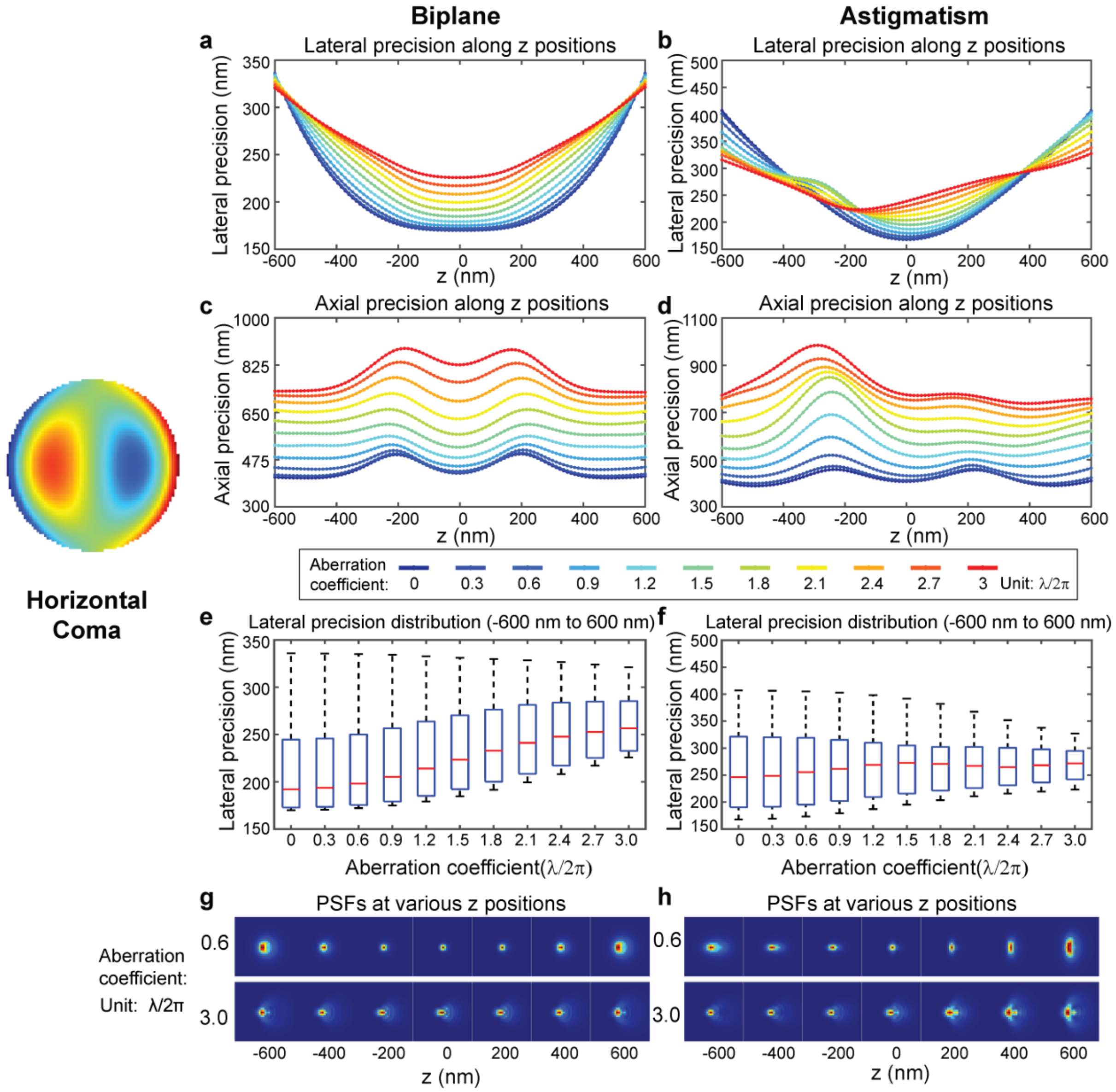

**Supplementary Fig. 4: Effects of horizontal coma on localization precision with varying amplitudes**. (a, b) Lateral localization precision along the z-axis with varying amplitudes of horizontal coma in biplane and astigmatism systems. The precision values are color-coded to represent aberration coefficients ranging from 0 to 3, with a step size of 0.3, measured in units of λ/2π. (c, d) Axial localization precision along the z-axis, similarly color-coded with varying horizontal coma amplitudes in biplane and astigmatism systems. (e, f) Distributions of lateral localization precision within the z range of -600 nm to 600 nm with varying amplitudes (ranging from 0 to 3) in biplane and astigmatism systems. Each boxplot displays the median, 25th, and 75th percentiles of the data, while whiskers extend to non-outlier extreme points. Outliers are individually marked with plus signs. Simulation conditions: $\lambda$ = 680 nm, $I$ = 0.5 photons/plane for the biplane system, $I$ = 1 photon for the astigmatism system, $bg$ = 0, $NA$ = 1.4, $n_{obj}$ = 1.52, pixel size = 130 nm, biplane distance = 400 nm, astigmatism value = 1.4 λ/2π. Lateral localization is calculated as the root mean square (RMS) of x- and y- localization precision. The region of interest (ROI) size was set to 32 × 32 pixels.

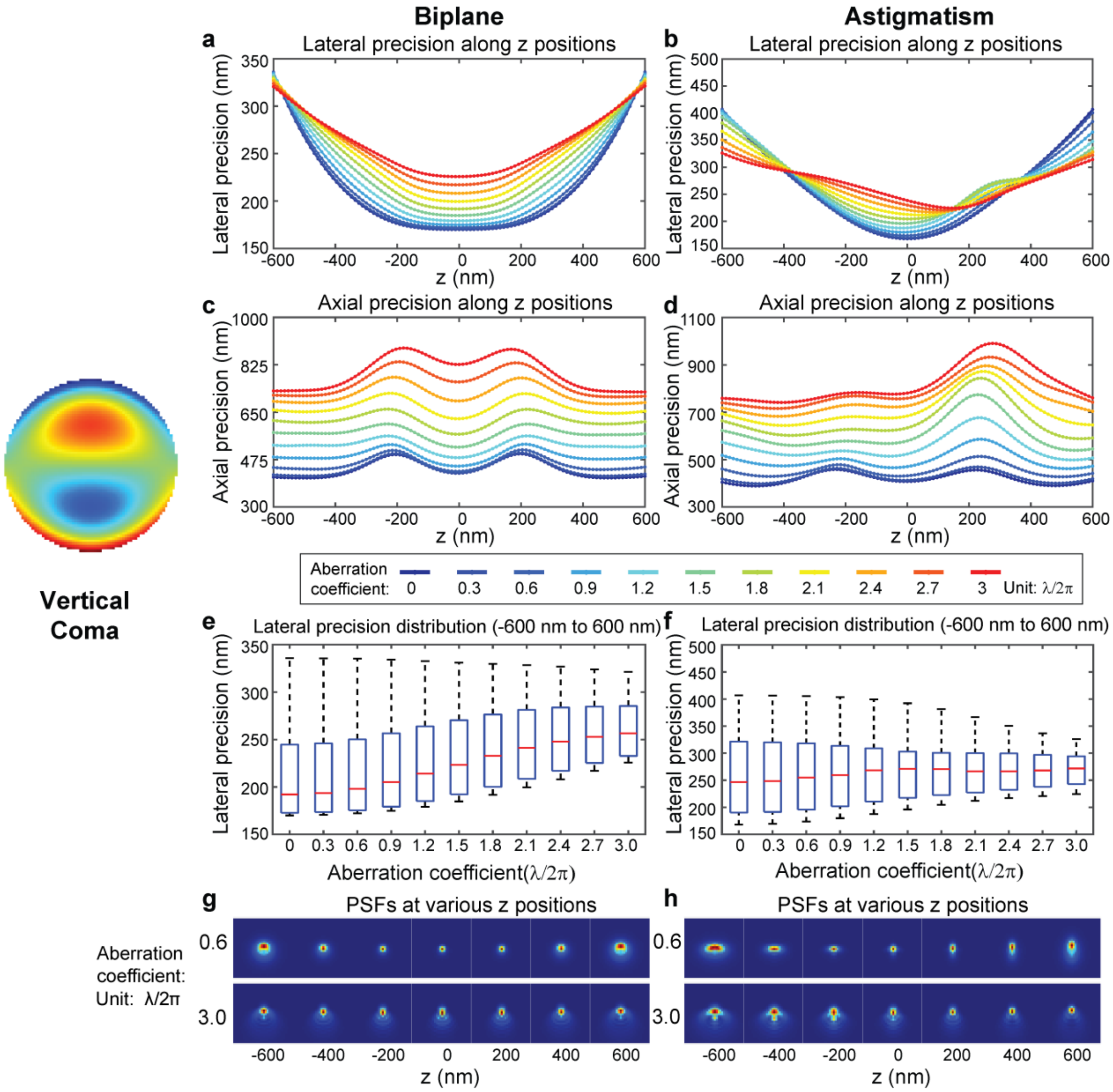

**Supplementary Fig. 5: Effects of vertical coma on localization precision with varying amplitudes**. (a, b) Lateral localization precision along the z-axis with varying amplitudes of vertical coma in biplane and astigmatism systems. The precision values are color-coded to represent aberration coefficients ranging from 0 to 3, with a step size of 0.3, measured in units of $\lambda/2\pi$. (c, d) Axial localization precision along the z-axis, similarly color-coded with varying vertical coma amplitudes in biplane and astigmatism systems. (e, f) Distributions of lateral localization precision within the z range of -600 nm to 600 nm with varying amplitudes (ranging from 0 to 3) in biplane and astigmatism systems. Each boxplot displays the median, 25th, and 75th percentiles of the data, while whiskers extend to non-outlier extreme points. Outliers are individually marked with plus signs. Simulation conditions $\lambda = 680$ nm, $I = 0.5$ photons/plane for the biplane system, $I = 1$ photon for the astigmatism system, $bg = 0$, $NA = 1.4$, $n_{obj} = 1.52$, pixel size = 130 nm, biplane distance = 400 nm, astigmatism value = 1.4 $\lambda/2\pi$. Lateral localization is calculated as the root mean square (RMS) of x- and y- localization precision. The region of interest (ROI) size was set to 32 × 32 pixels.

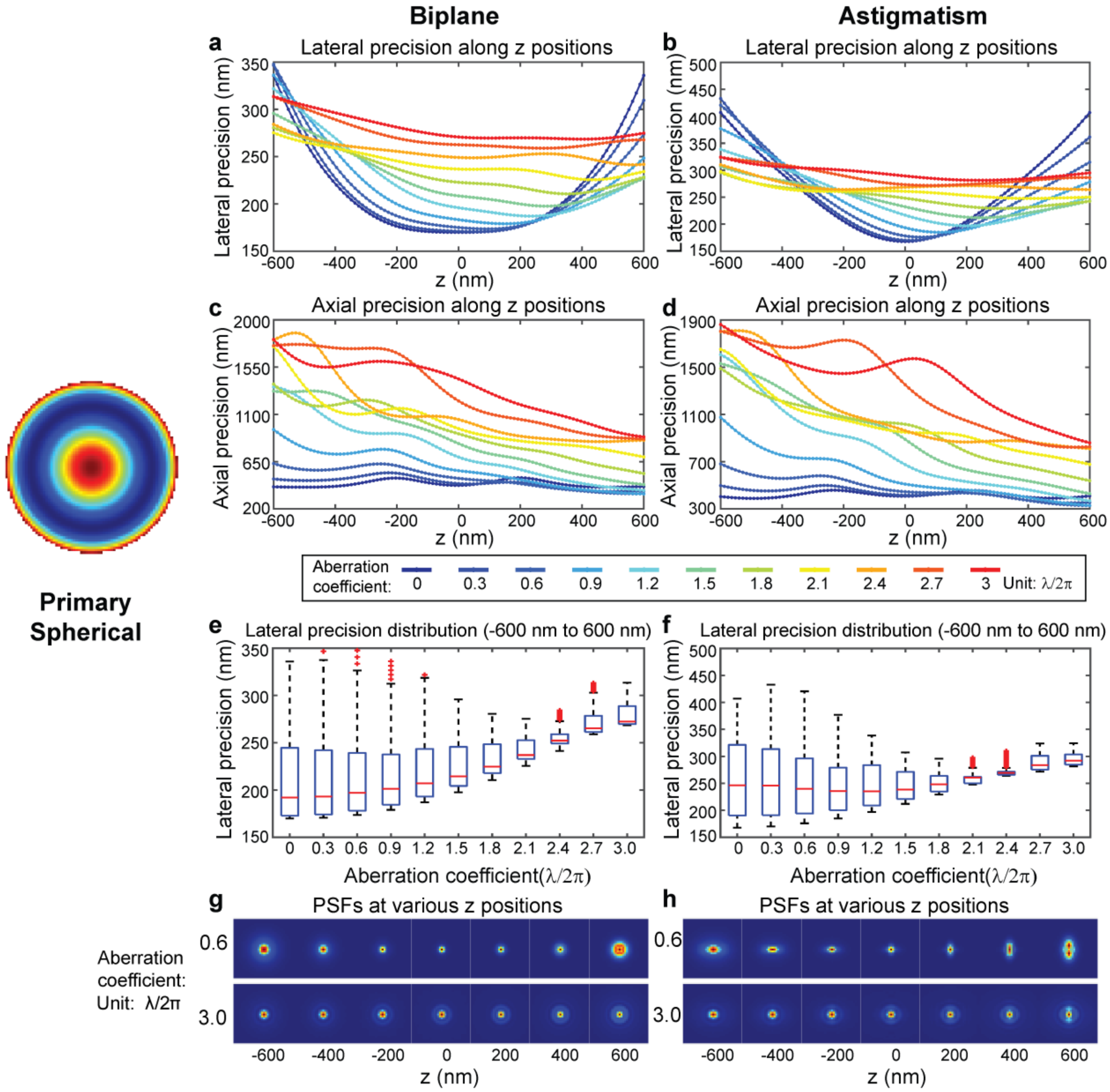

**Supplementary Fig. 6: Effects of primary spherical on localization precision with varying amplitudes**. (a, b) Lateral localization precision along the z-axis with varying amplitudes of primary spherical in biplane and astigmatism systems. The precision values are color-coded to represent aberration coefficients ranging from 0 to 3, with a step size of 0.3, measured in units of λ/2π. (c, d) Axial localization precision along the z-axis, similarly color-coded with varying primary spherical amplitudes in biplane and astigmatism systems. (e, f) Distributions of lateral localization precision within the z range of -600 nm to 600 nm with varying amplitudes (ranging from 0 to 3) in biplane and astigmatism systems. Each boxplot displays the median, 25th, and 75th percentiles of the data, while whiskers extend to non-outlier extreme points. Outliers are individually marked with plus signs. Simulation conditions: $\lambda$ = 680 nm, $I$ = 0.5 photons/plane for the biplane system, $I$ = 1 photon for the astigmatism system, $bg$ = 0, $NA$ = 1.4, $n_{obj}$ = 1.52, pixel size = 130 nm, biplane distance = 400 nm, astigmatism value = 1.4 λ/2π. Lateral localization is calculated as the root mean square (RMS) of x- and y- localization precision. The region of interest (ROI) size was set to 32 × 32 pixels.

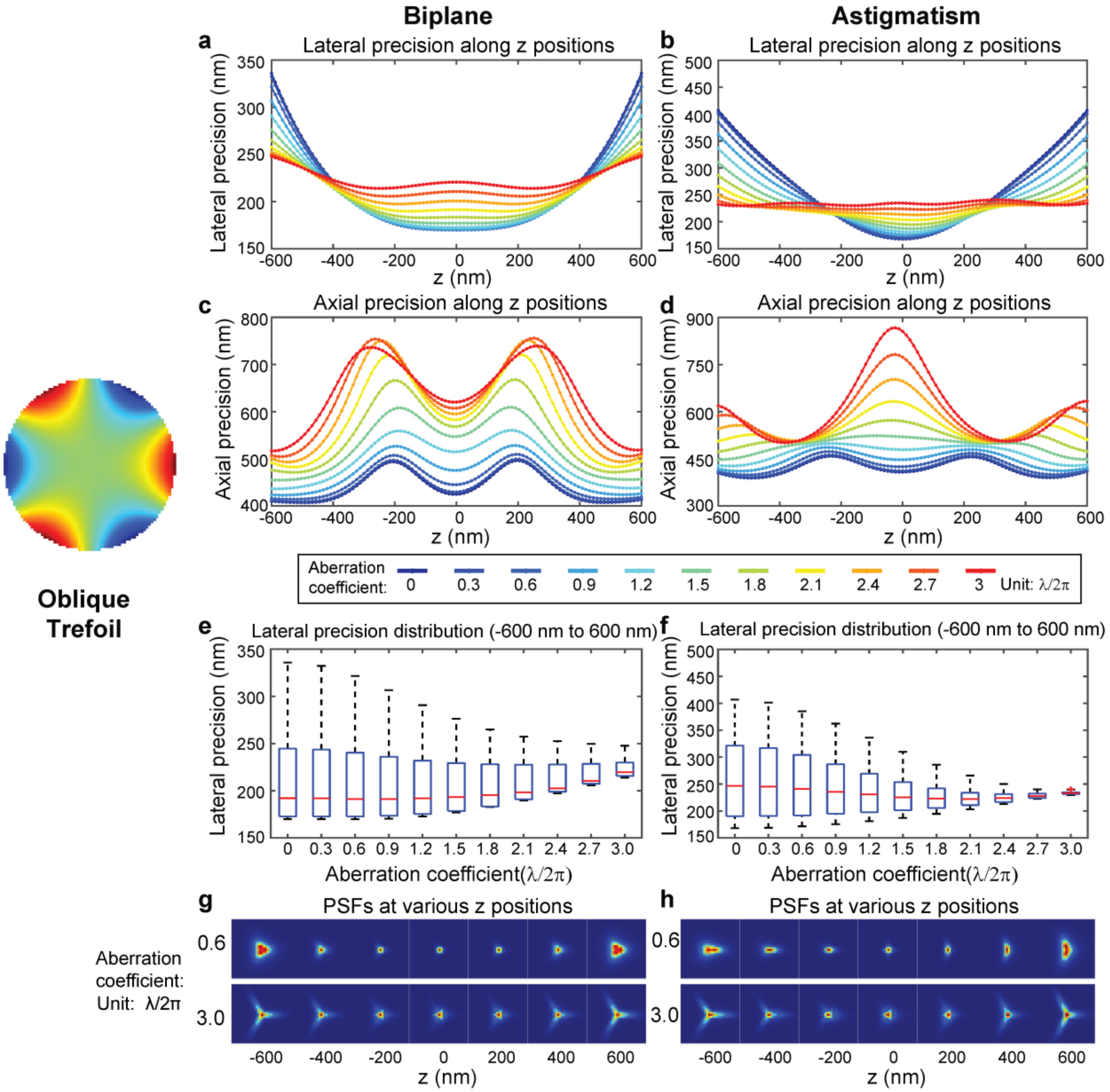

**Supplementary Fig. 7: Effects of oblique trefoil on localization precision with varying amplitudes**. (a, b) Lateral localization precision along the z-axis with varying amplitudes of oblique trefoil in biplane and astigmatism systems. The precision values are color-coded to represent aberration coefficients ranging from 0 to 3, with a step size of 0.3, measured in units of $\lambda/2\pi$. (c, d) Axial localization precision along the z-axis, similarly color-coded with varying oblique trefoil amplitudes in biplane and astigmatism systems. (e, f) Distributions of lateral localization precision within the z range of -600 nm to 600 nm with varying amplitudes (ranging from 0 to 3) in biplane and astigmatism systems. Each boxplot displays the median, 25th, and 75th percentiles of the data, while whiskers extend to non-outlier extreme points. Outliers are individually marked with plus signs. Simulation conditions: $\lambda$ = 680 nm, $I$ = 0.5 photons/plane for the biplane system, $I$ = 1 photon for the astigmatism system, $bg$ = 0, $NA$ = 1.4, $n_{obj}$ = 1.52, pixel size = 130 nm, biplane distance = 400 nm, astigmatism value = 1.4 $\lambda/2\pi$. Lateral localization is calculated as the root mean square (RMS) of x- and y- localization precision. The region of interest (ROI) size was set to 32 × 32 pixels.

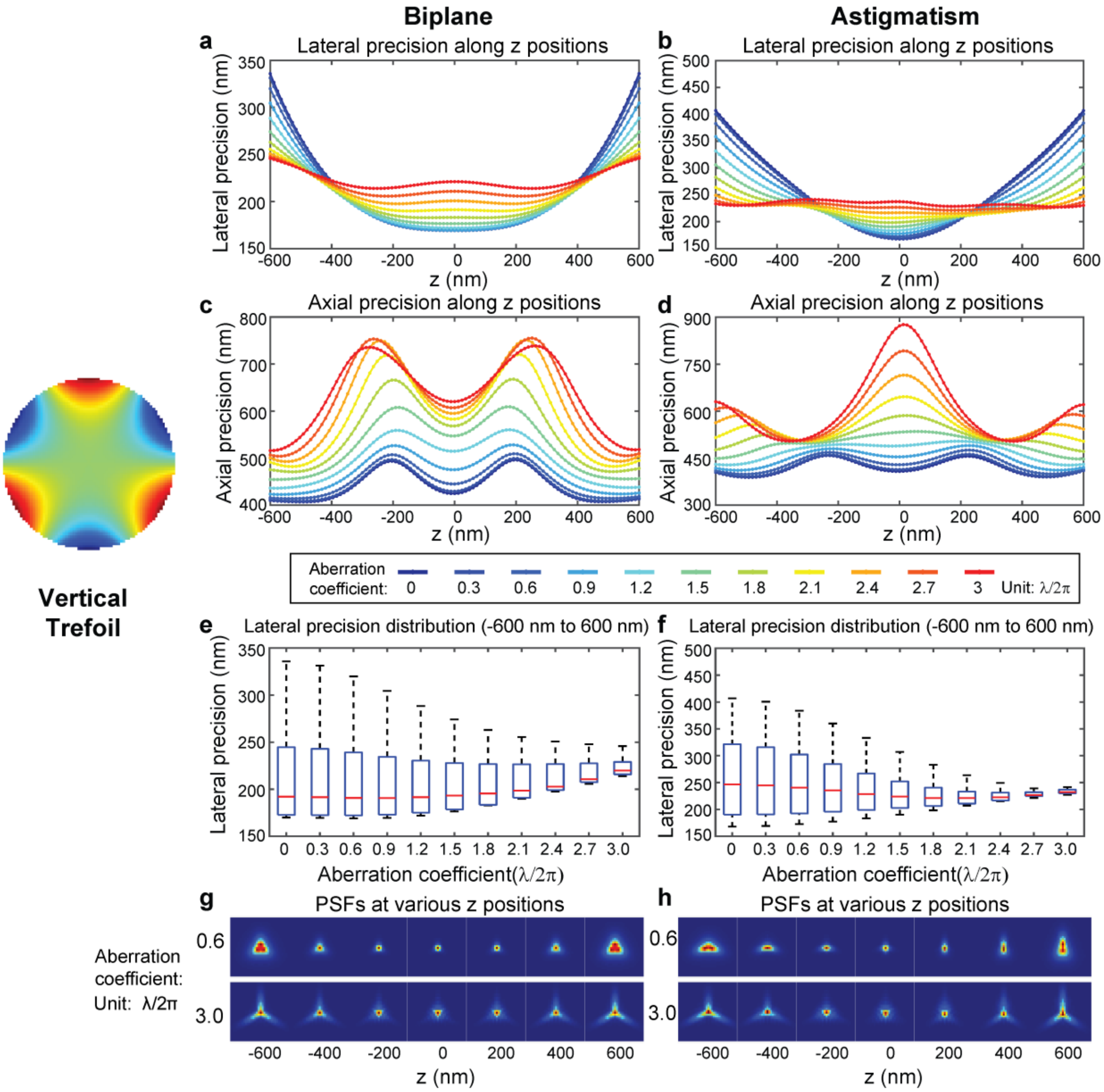

**Supplementary Fig. 8: Effects of vertical trefoil on localization precision with varying amplitudes**. (a, b) Lateral localization precision along the z-axis with varying amplitudes of vertical trefoil in biplane and astigmatism systems. The precision values are color-coded to represent aberration coefficients ranging from 0 to 3, with a step size of 0.3, measured in units of λ/2π. (c, d) Axial localization precision along the z-axis, similarly color-coded with varying vertical trefoil amplitudes in biplane and astigmatism systems. (e, f) Distributions of lateral localization precision within the z range of -600 nm to 600 nm with varying amplitudes (ranging from 0 to 3) in biplane and astigmatism systems. Each boxplot displays the median, 25th, and 75th percentiles of the data, while whiskers extend to non-outlier extreme points. Outliers are individually marked with plus signs. Simulation conditions: $\lambda$ = 680 nm, $I$ = 0.5 photons/plane for the biplane system, $I$ = 1 photon for the astigmatism system, $bg$ = 0, $NA$ = 1.4, $n_{obj}$ = 1.52, pixel size = 130 nm, biplane distance = 400 nm, astigmatism value = 1.4 λ/2π. Lateral localization is calculated as the root mean square (RMS) of x- and y- localization precision. The region of interest (ROI) size was set to 32 × 32 pixels.

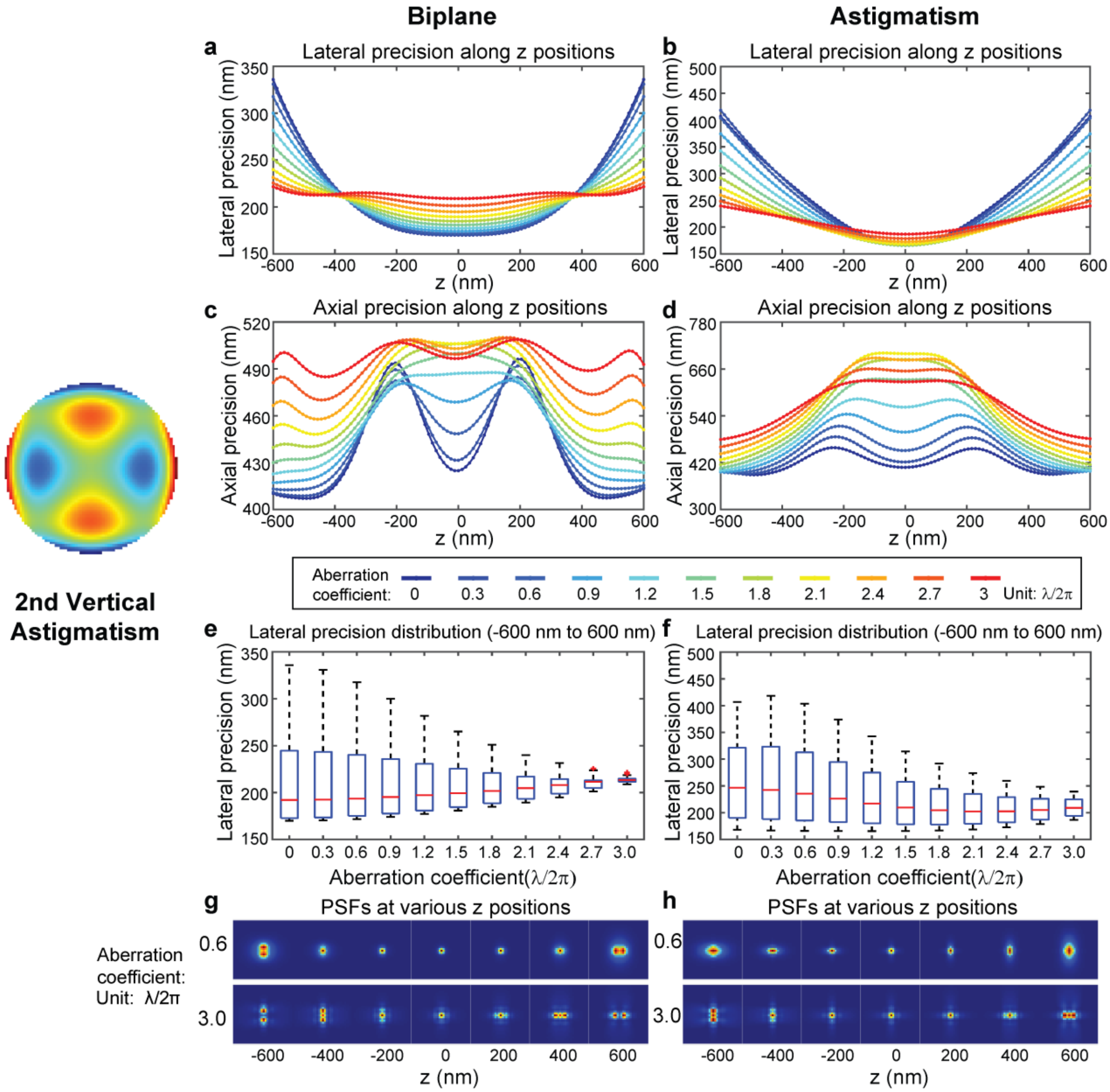

**Supplementary Fig. 9: Effects of secondary vertical astigmatism on localization precision with varying amplitudes**. (a, b) Lateral localization precision along the z-axis with varying amplitudes of secondary vertical astigmatism in biplane and astigmatism systems. The precision values are color-coded to represent aberration coefficients ranging from 0 to 3, with a step size of 0.3, measured in units of λ/2π. (c, d) Axial localization precision along the z-axis, similarly color-coded with varying secondary vertical astigmatism amplitudes in biplane and astigmatism systems. (e, f) Distributions of lateral localization precision within the z range of -600 nm to 600 nm with varying amplitudes (ranging from 0 to 3) in biplane and astigmatism systems. Each boxplot displays the median, 25th, and 75th percentiles of the data, while whiskers extend to non-outlier extreme points. Outliers are individually marked with plus signs. Simulation conditions: $\lambda$ = 680 nm, $I$ = 0.5 photons/plane for the biplane system, $I$ = 1 photon for the astigmatism system, $bg$ = 0, $NA$ = 1.4, $n_{obj}$ = 1.52, pixel size = 130 nm, biplane distance = 400 nm, astigmatism value = 1.4 λ/2π. Lateral localization is calculated as the root mean square (RMS) of x- and y- localization precision. The region of interest (ROI) size was set to 32 × 32 pixels.

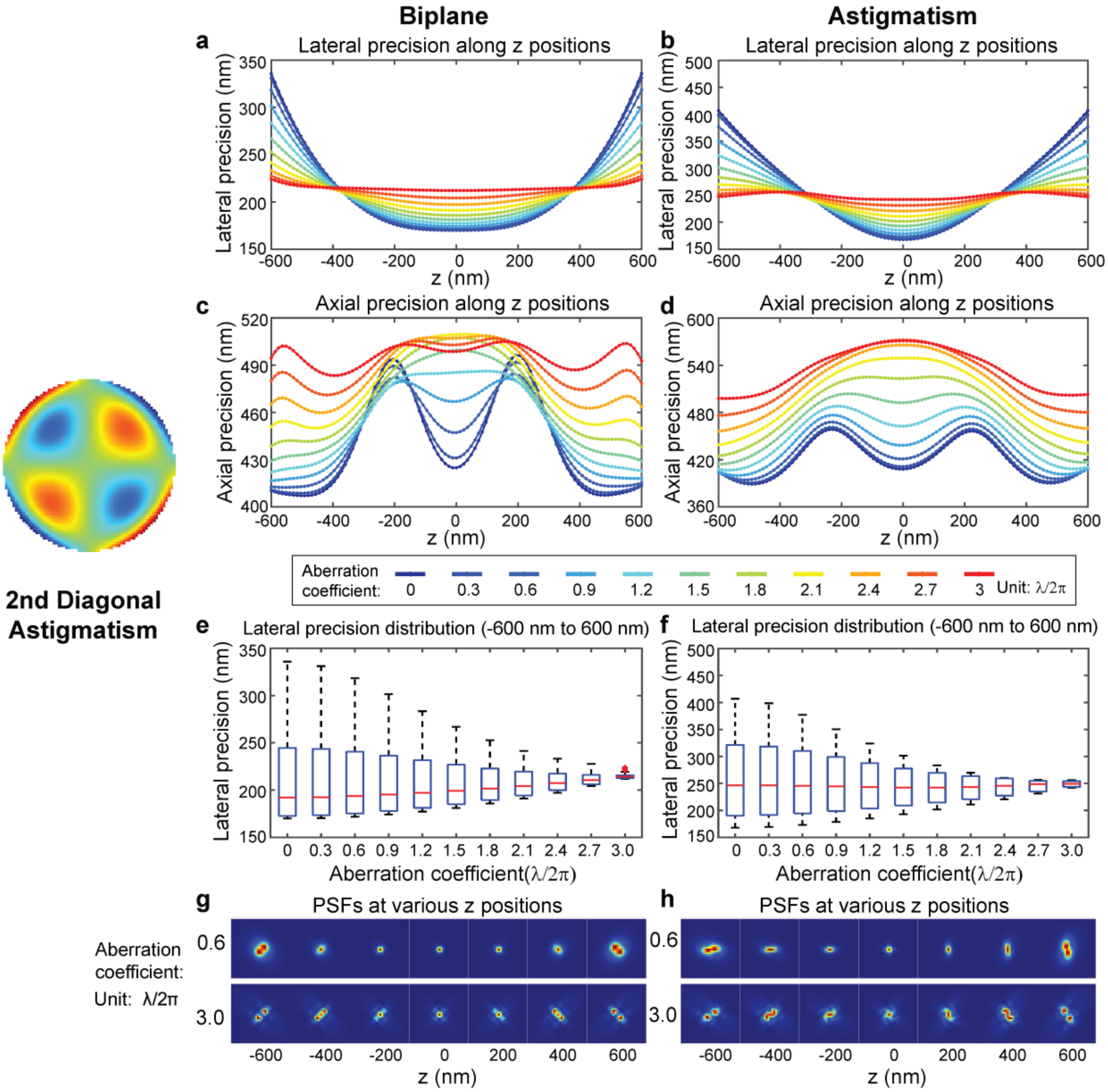

**Supplementary Fig. 10: Effects of secondary diagonal astigmatism on localization precision with varying amplitudes**. (a, b) Lateral localization precision along the z-axis with varying amplitudes of secondary diagonal astigmatism in biplane and astigmatism systems. The precision values are color-coded to represent aberration coefficients ranging from 0 to 3, with a step size of 0.3, measured in units of λ/2π. (c, d) Axial localization precision along the z-axis, similarly color-coded with varying secondary diagonal astigmatism amplitudes in biplane and astigmatism systems. (e, f) Distributions of lateral localization precision within the z range of -600 nm to 600 nm with varying amplitudes (ranging from 0 to 3) in biplane and astigmatism systems. Each boxplot displays the median, 25th, and 75th percentiles of the data, while whiskers extend to non-outlier extreme points. Outliers are individually marked with plus signs. Simulation conditions: $\lambda$ = 680 nm, $I$ = 0.5 photons/plane for the biplane system, $I$ = 1 photon for the astigmatism system, $bg$ = 0, $NA$ = 1.4, $n_{obj}$ = 1.52, pixel size = 130 nm, biplane distance = 400 nm, astigmatism value = 1.4 λ/2π. Lateral localization is calculated as the root mean square (RMS) of x- and y- localization precision. The region of interest (ROI) size was set to 32 × 32 pixels.

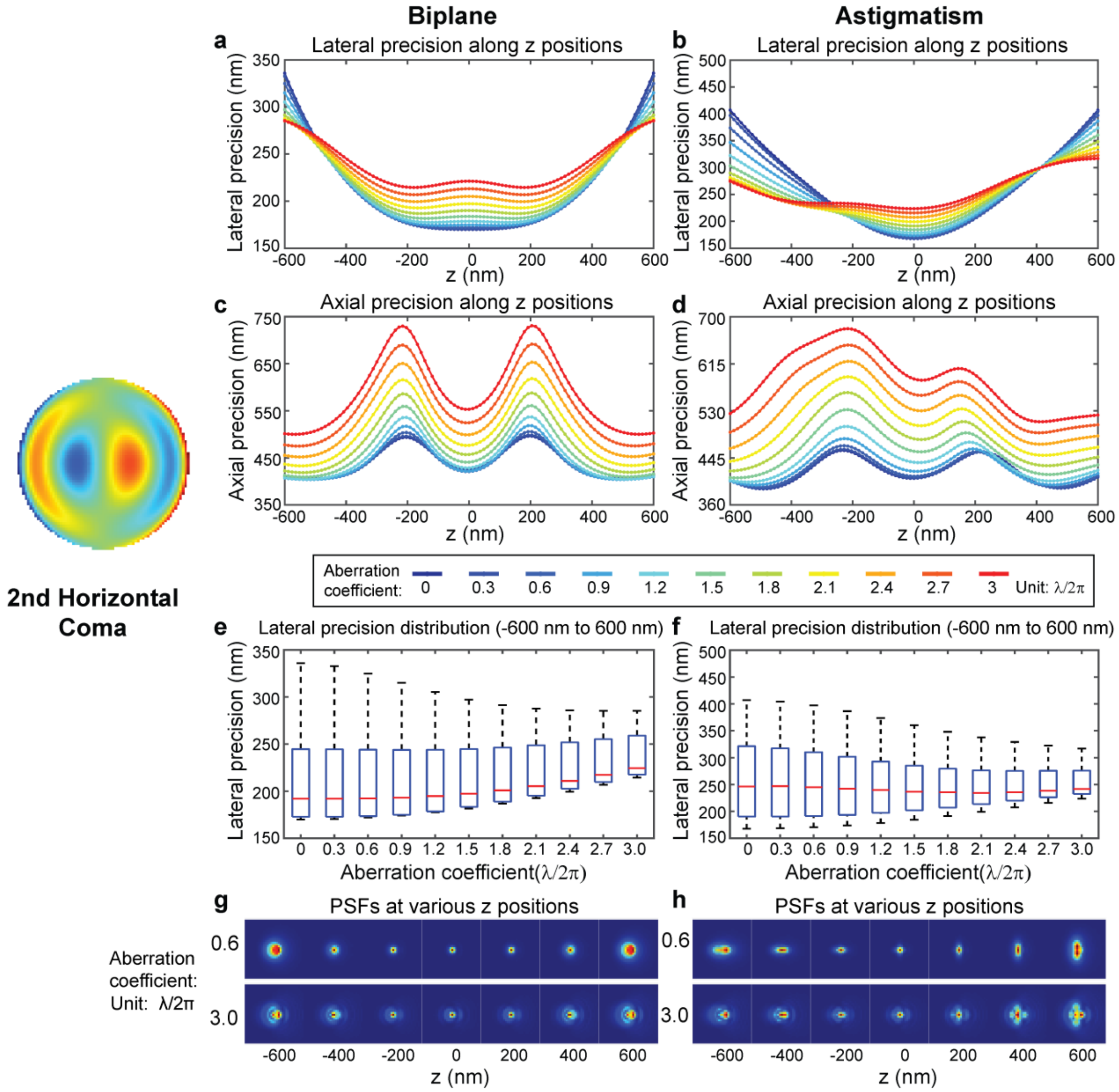

**Supplementary Fig. 11: Effects of secondary horizontal coma on localization precision with varying amplitudes**. (a, b) Lateral localization precision along the z-axis with varying amplitudes of secondary horizontal coma in biplane and astigmatism systems. The precision values are color-coded to represent aberration coefficients ranging from 0 to 3, with a step size of 0.3, measured in units of λ/2π. (c, d) Axial localization precision along the z-axis, similarly color-coded with varying secondary horizontal coma amplitudes in biplane and astigmatism systems. (e, f) Distributions of lateral localization precision within the z range of -600 nm to 600 nm with varying amplitudes (ranging from 0 to 3) in biplane and astigmatism systems. Each boxplot displays the median, 25th, and 75th percentiles of the data, while whiskers extend to non-outlier extreme points. Outliers are individually marked with plus signs. Simulation conditions: $\lambda$ = 680 nm, $I$ = 0.5 photons/plane for the biplane system, $I$ = 1 photon for the astigmatism system, $bg$ = 0, $NA$ = 1.4, $n_{obj}$ = 1.52, pixel size = 130 nm, biplane distance = 400 nm, astigmatism value = 1.4 λ/2π. Lateral localization is calculated as the root mean square (RMS) of x- and y- localization precision. The region of interest (ROI) size was set to 32 × 32 pixels.

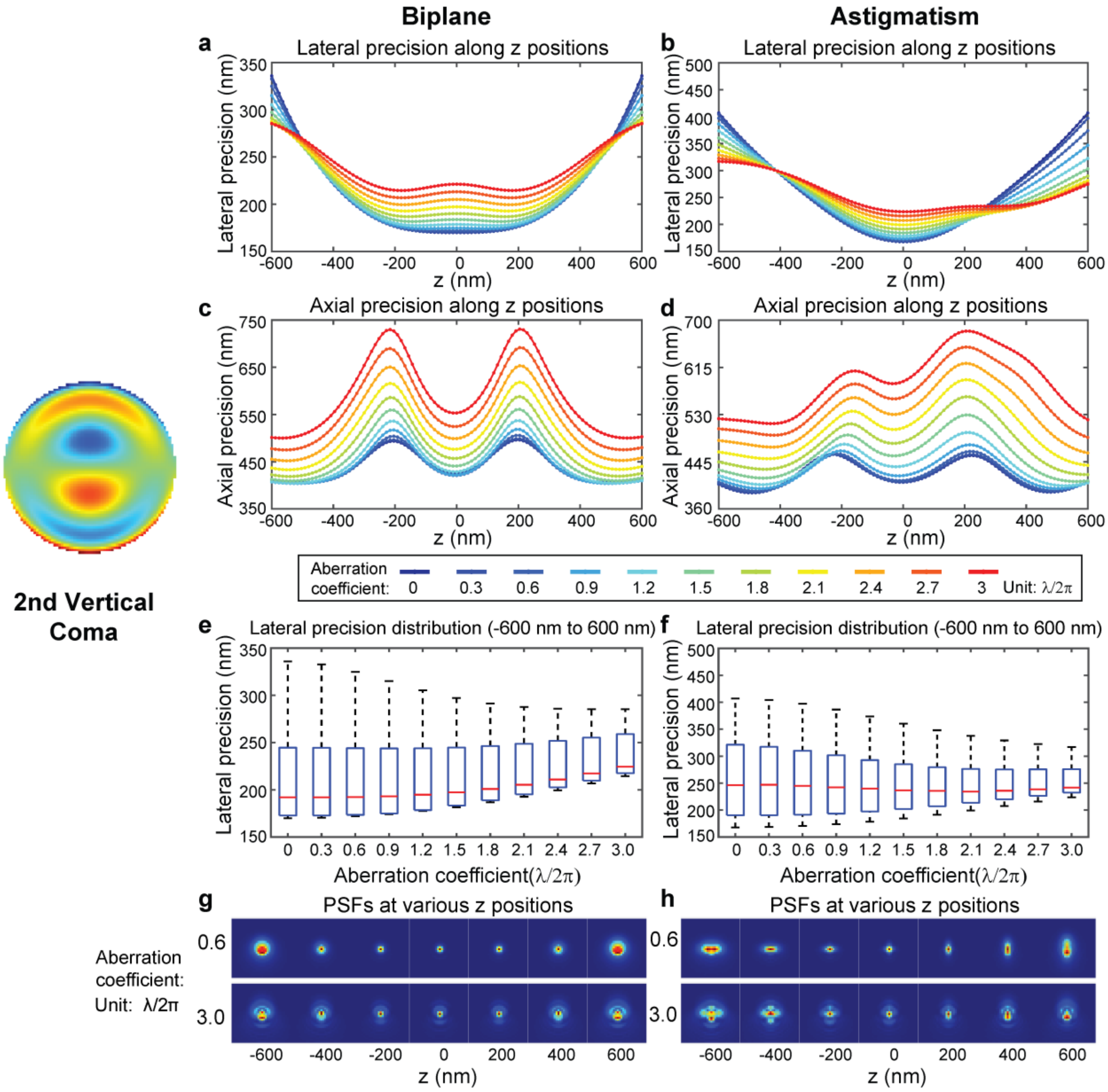

**Supplementary Fig. 12: Effects of secondary vertical coma on localization precision with varying amplitudes**. (a, b) Lateral localization precision along the z-axis with varying amplitudes of secondary vertical coma in biplane and astigmatism systems. The precision values are color-coded to represent aberration coefficients ranging from 0 to 3, with a step size of 0.3, measured in units of λ/2π. (c, d) Axial localization precision along the z-axis, similarly color-coded with varying secondary vertical coma amplitudes in biplane and astigmatism systems. (e, f) Distributions of lateral localization precision within the z range of -600 nm to 600 nm with varying amplitudes (ranging from 0 to 3) in biplane and astigmatism systems. Each boxplot displays the median, 25th, and 75th percentiles of the data, while whiskers extend to non-outlier extreme points. Outliers are individually marked with plus signs. Simulation conditions: $\lambda$ = 680 nm, $I$ = 0.5 photons/plane for the biplane system, $I$ = 1 photon for the astigmatism system, $bg$ = 0, $NA$ = 1.4, $n_{obj}$ = 1.52, pixel size = 130 nm, biplane distance = 400 nm, astigmatism value = 1.4 λ/2π. Lateral localization is calculated as the root mean square (RMS) of x- and y- localization precision. The region of interest (ROI) size was set to 32 × 32 pixels.

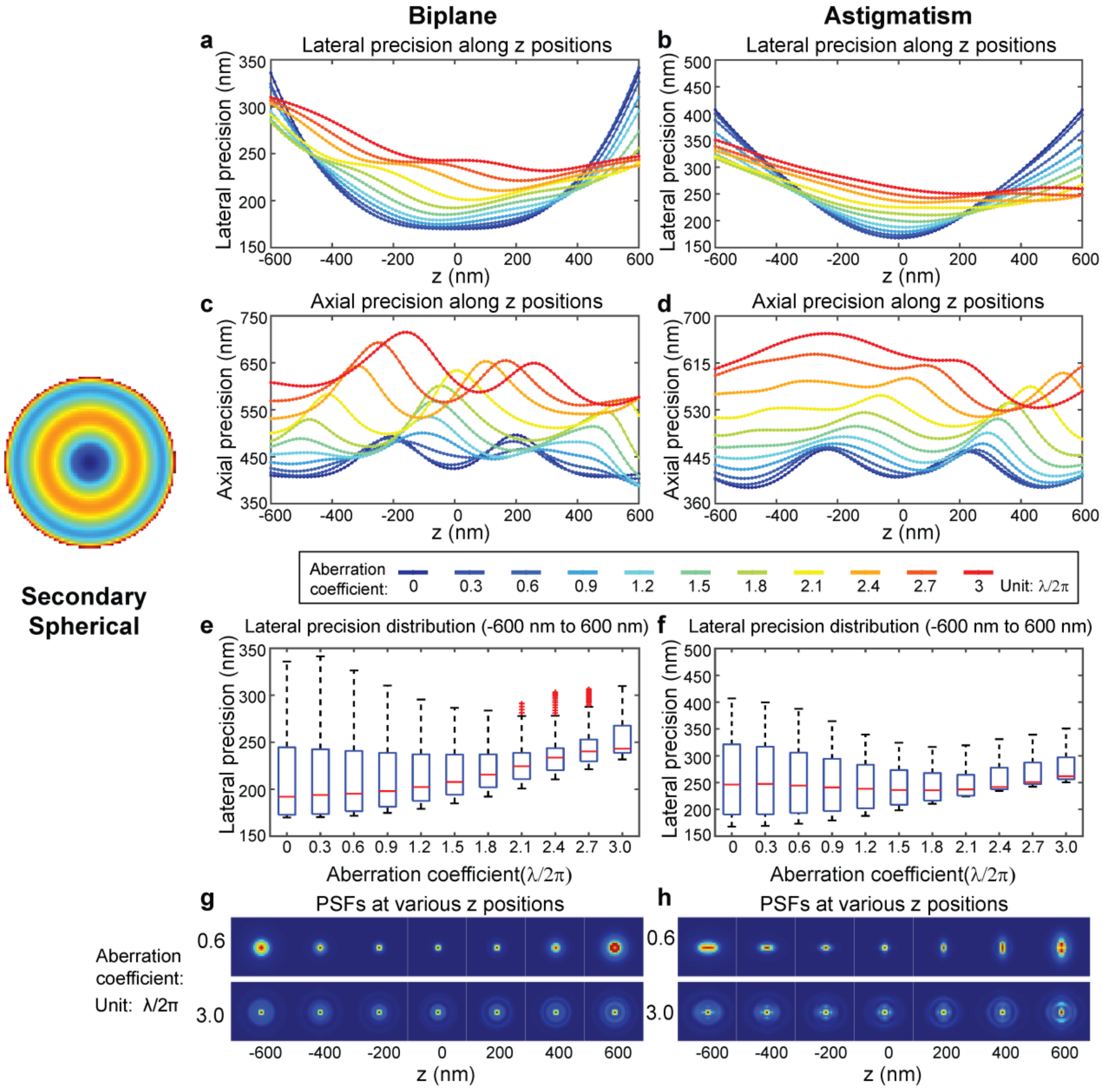

**Supplementary Fig. 13: Effects of secondary spherical on localization precision with varying amplitudes**. (a, b) Lateral localization precision along the z-axis with varying amplitudes of secondary spherical in biplane and astigmatism systems. The precision values are color-coded to represent aberration coefficients ranging from 0 to 3, with a step size of 0.3, measured in units of λ/2π. (c, d) Axial localization precision along the z-axis, similarly color-coded with varying secondary spherical amplitudes in biplane and astigmatism systems. (e, f) Distributions of lateral localization precision within the z range of -600 nm to 600 nm with varying amplitudes (ranging from 0 to 3) in biplane and astigmatism systems. Each boxplot displays the median, 25th, and 75th percentiles of the data, while whiskers extend to non-outlier extreme points. Outliers are individually marked with plus signs. Simulation conditions: $\lambda$ = 680 nm, $I$ = 0.5 photons/plane for the biplane system, $I$ = 1 photon for the astigmatism system, $bg$ = 0, $NA$ = 1.4, $n_{obj}$ = 1.52, pixel size = 130 nm, biplane distance = 400 nm, astigmatism value = 1.4 λ/2π. Lateral localization is calculated as the root mean square (RMS) of x- and y- localization precision. The region of interest (ROI) size was set to 32 × 32 pixels.

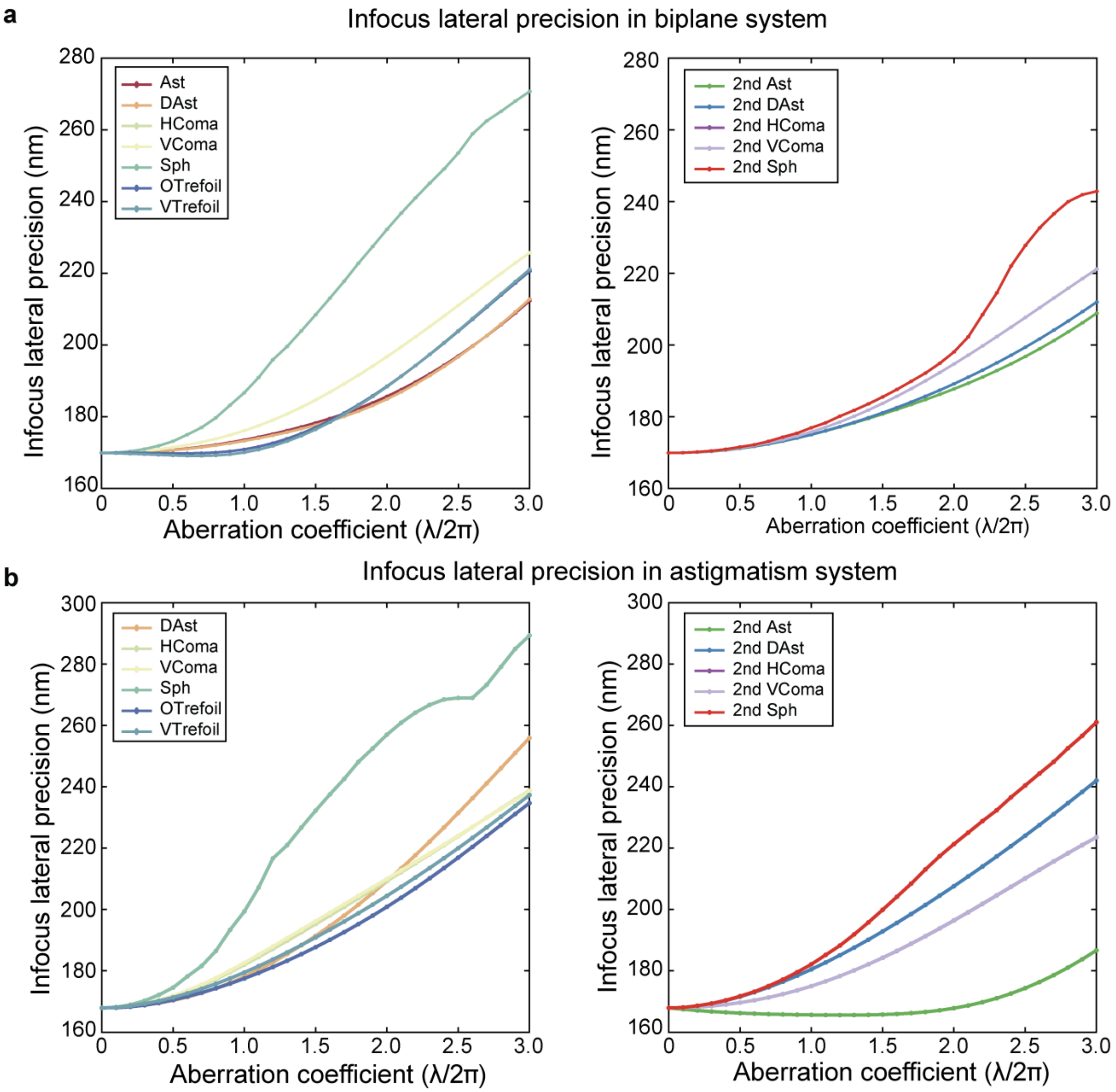

**Supplementary Fig. 14: In-focus (at z = 0) lateral localization precision with varying aberration types and amplitudes in (a) biplane and (b) astigmatism systems.** Simulation conditions: $\lambda$ = 680 nm, $I$ = 0.5 photons/plane for the biplane system, $I$ = 1 photon for the astigmatism system, $bg$ = 0, $NA$ = 1.4, $n_{obj}$ = 1.52, pixel size = 130 nm, biplane distance = 400 nm, astigmatism value = 1.4 λ/2π. Lateral localization is calculated as the root mean square (RMS) of x- and y- localization precision. The region of interest (ROI) size was set to 32 × 32 pixels. Abbreviations: Ast (vertical astigmatism), DAst (diagonal astigmatism), HComa (horizontal coma), VComa (vertical coma), Sph (primary spherical), OTrefoil (oblique trefoil), VTrefoil (vertical trefoil), 2nd Ast (secondary vertical astigmatism), 2nd DAst (secondary diagonal astigmatism), 2nd HComa (secondary horizontal coma), 2nd VComa (secondary vertical coma), 2nd Sph (secondary spherical).

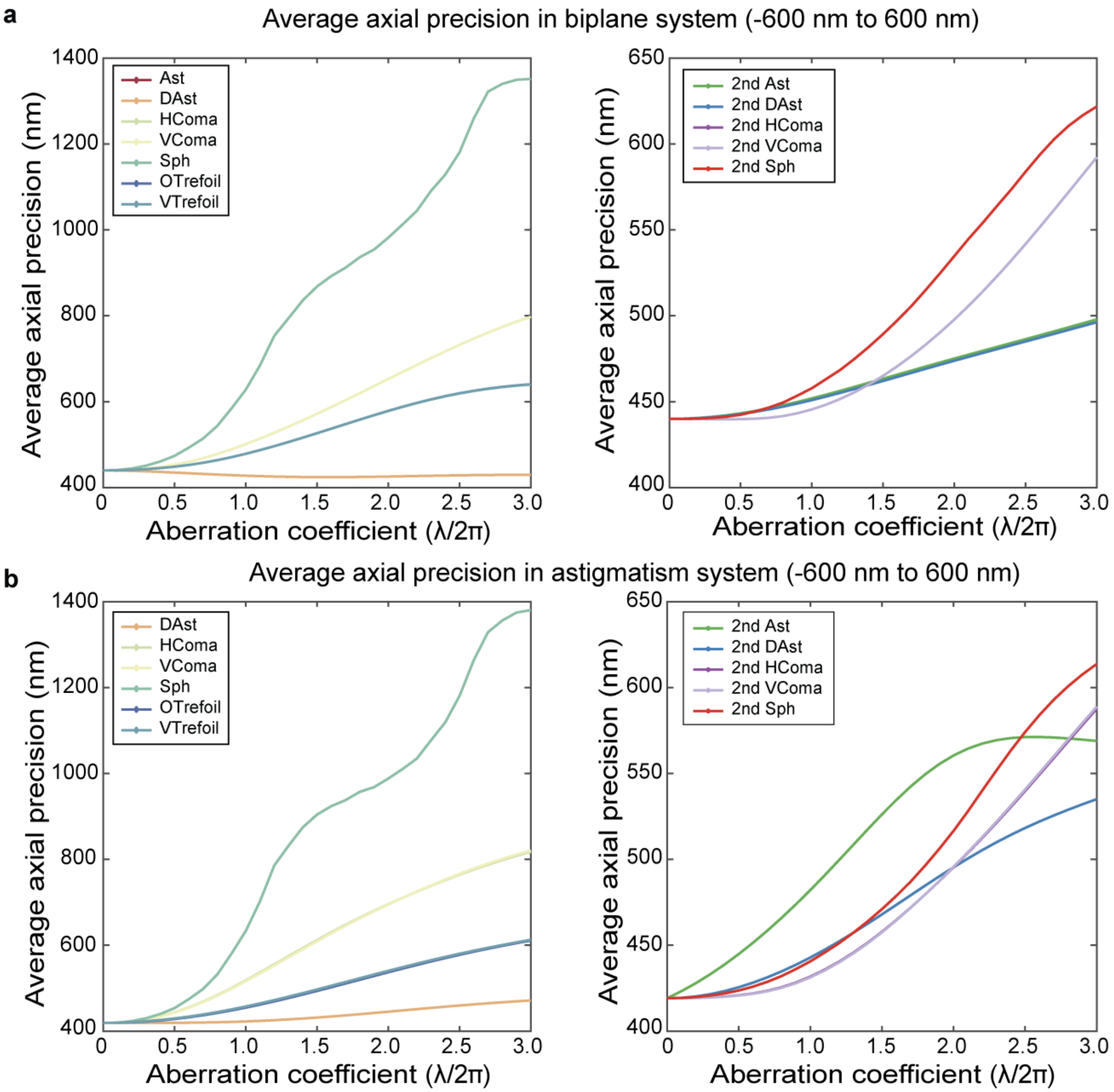

**Supplementary Fig. 15: Average axial localization precision with varying aberration types and amplitudes in (a) biplane and (b) astigmatism systems.** The average is calculated from 101 frames within the -600 nm to 600 nm z-range with equal spacing. Simulation conditions: $\lambda$ = 680 nm, $I$ = 0.5 photons/plane for the biplane system, $I$ = 1 photon for the astigmatism system, $bg$ = 0, $NA$ = 1.4, $n_{obj}$ = 1.52, pixel size = 130 nm, biplane distance = 400 nm, astigmatism value = 1.4 λ/2π. The region of interest (ROI) size was set to 32 × 32 pixels. Abbreviations: Ast (vertical astigmatism), DAst (diagonal astigmatism), HComa (horizontal coma), VComa (vertical coma), Sph (primary spherical), OTrefoil (oblique trefoil), VTrefoil (vertical trefoil), 2nd Ast (secondary vertical astigmatism), 2nd DAst (secondary diagonal astigmatism), 2nd HComa (secondary horizontal coma), 2nd VComa (secondary vertical coma), 2nd Sph (secondary spherical).

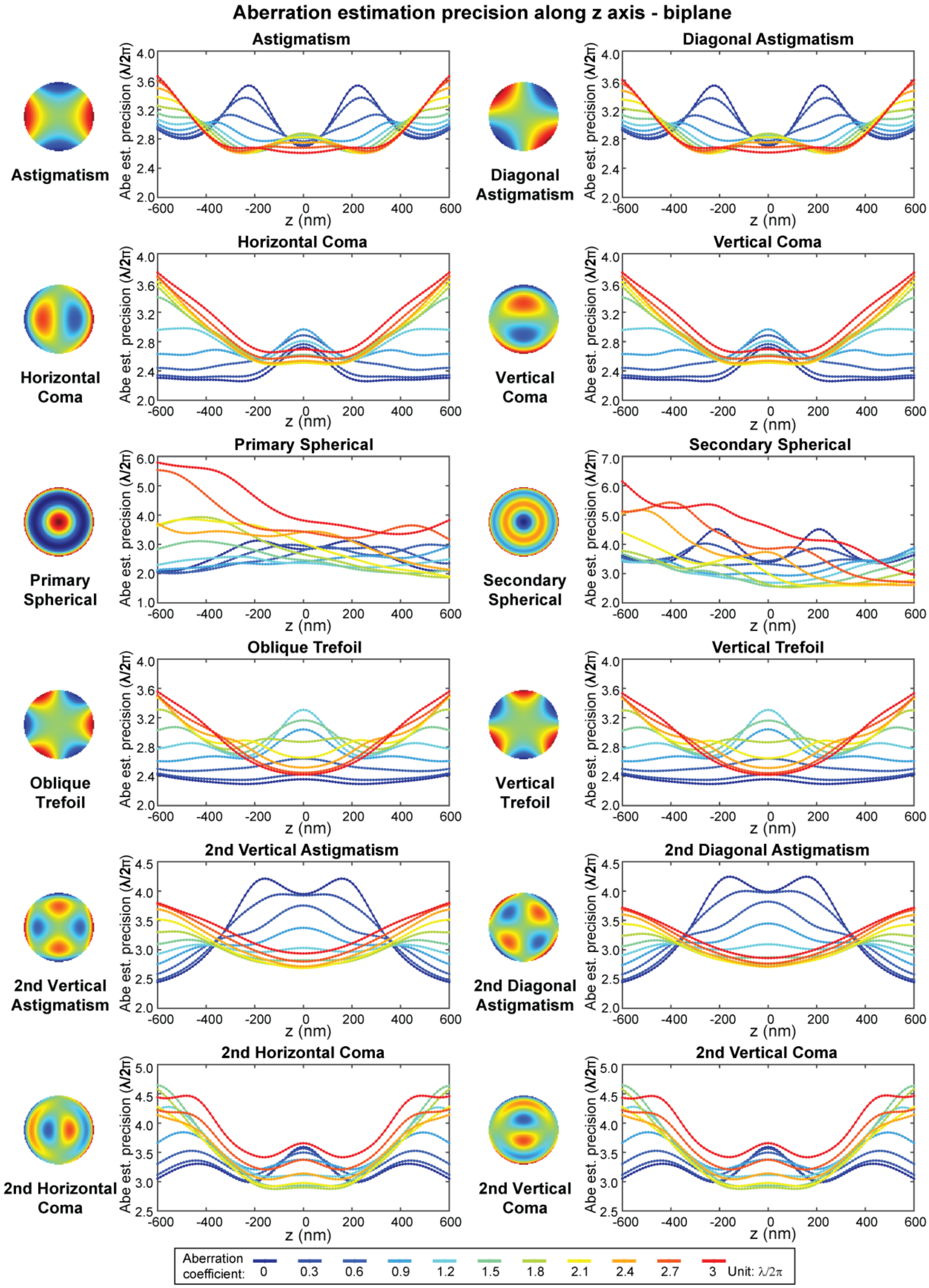
Aberration estimation precision along z axis - biplane
Astigmatism
Diagonal Astigmatism
Horizontal Coma
Vertical Coma
Primary Spherical
Secondary Spherical
Oblique Trefoil
Vertical Trefoil
2nd Vertical Astigmatism
2nd Diagonal Astigmatism
2nd Horizontal Coma
2nd Vertical Coma
Abe est. precision (λ/2π)
z (nm)
Aberration coefficient: 0 0.3 0.6 0.9 1.2 1.5 1.8 2.1 2.4 2.7 3 Unit: λ/2π

**Supplementary Fig. 16: Aberration estimation precision at different z positions with varying aberration types and amplitudes in biplane system.** Each panel represents the precision of a specific type of aberration with varying amplitudes within the system, ranging from 0 to 3, with a step size of 0.3, measured in units of $\lambda/2\pi$. Simulation conditions: $\lambda$ = 680 nm, $I$ = 0.5 photons/plane for the biplane system, $bg$ = 0, $NA$ = 1.4, $n_{obj}$ = 1.52, pixel size = 130 nm, biplane distance = 400 nm. The region of interest (ROI) size was set to 32 × 32 pixels.

**Aberration estimation precision along z axis - astigmatism**

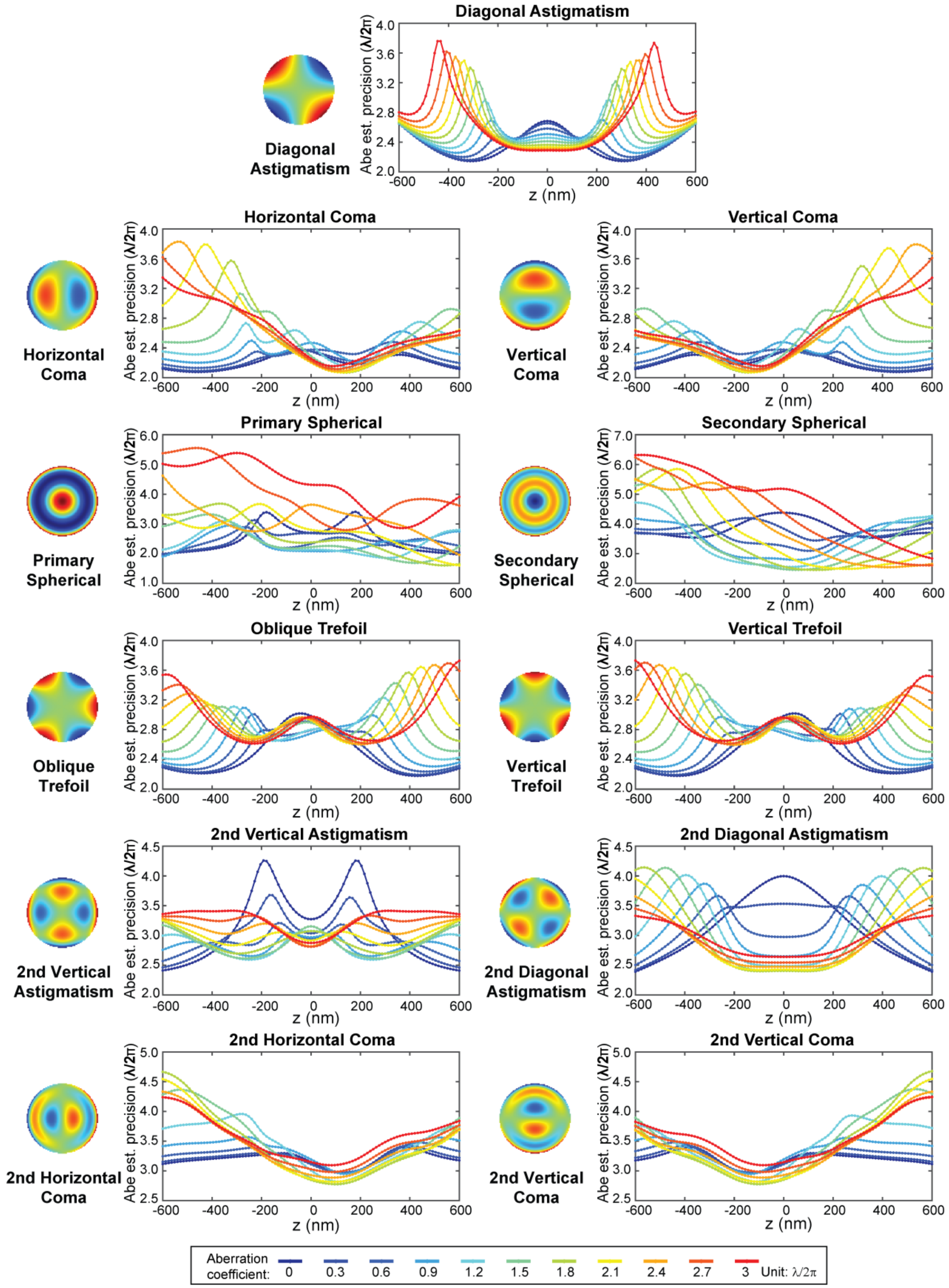

**Supplementary Fig. 17: Aberration estimation precision at different z positions with varying aberration types and amplitudes in astigmatism system.** Each panel represents the precision of a specific type of aberration with varying amplitudes within the system, ranging from 0 to 3, with a step size of 0.3, measured in units of $\lambda/2\pi$. Simulation conditions: $\lambda$ = 680 nm, $I$ = 1 photon for the astigmatism system, $bg$ = 0, $NA$ = 1.4, $n_{obj}$ = 1.52, pixel size = 130 nm, astigmatism value = 1.4 $\lambda/2\pi$. The region of interest (ROI) size was set to 32 × 32 pixels.

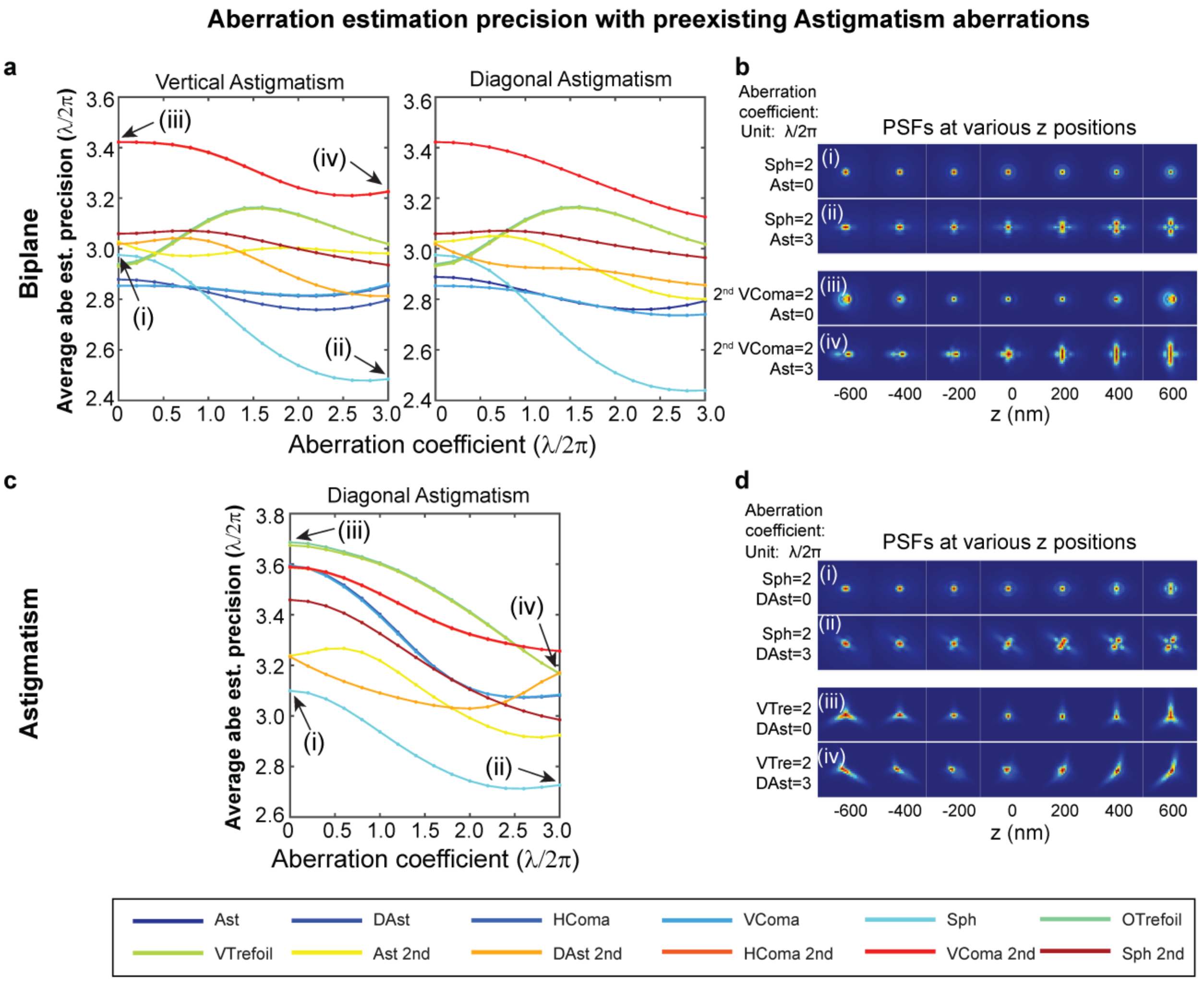

**Supplementary Fig. 18: Aberration estimation precision with preexisting astigmatism aberrations.** (a, c) Precision of aberration estimation when estimating each aberration at an amplitude of 2 λ/2π while introducing varying levels of vertical (left) and diagonal (right) astigmatism in biplane (a) and diagonal astigmatism in astigmatism system (c). The precision values are calculated by averaging within the -600 nm to 600 nm z range. The rainbow colors represent different aberration modes, ranging from vertical astigmatism to secondary spherical aberration. (b, d) Examples of PSFs within the -600 nm to 600 nm range, corresponding to cases as indicated in the figures. Simulation conditions: $\lambda$ = 680 nm, $I$ = 0.5 photons/plane for the biplane system, $I$ = 1 photon for the astigmatism system, $bg$ = 0, $NA$ = 1.4, $n_{obj}$ = 1.52, pixel size = 130 nm, biplane distance = 400 nm, astigmatism value = 1.4 λ/2π. The region of interest (ROI) size was set to 32 × 32 pixels.

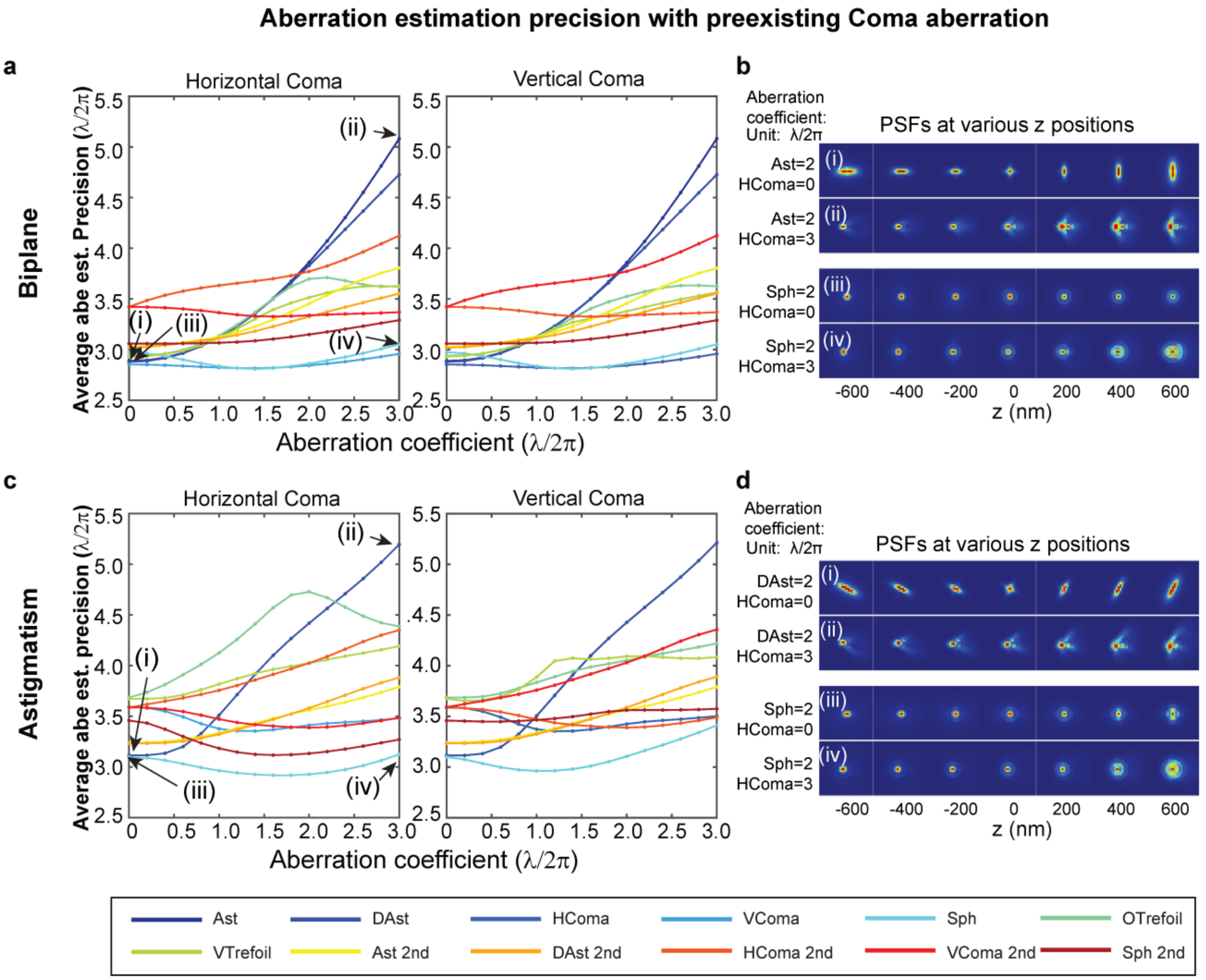

**Supplementary Fig. 19: Aberration estimation precision with preexisting coma aberrations.** (a, c) Precision of aberration estimation when estimating each aberration at an amplitude of 2 λ/2π while introducing varying levels of horizontal (left) and vertical (right) coma in (a) biplane and (c) astigmatism system. The precision values are calculated by averaging within the -600 nm to 600 nm z range. The rainbow colors represent different aberration modes, ranging from vertical astigmatism to secondary spherical aberration. (b, d) Examples of PSFs within the -600 nm to 600 nm range, corresponding to cases as indicated in the figures. Simulation conditions: $\lambda$ = 680 nm, $I$ = 0.5 photons/plane for the biplane system, $I$ = 1 photon for the astigmatism system, $bg$ = 0, $NA$ = 1.4, $n_{obj}$ = 1.52, pixel size = 130 nm, biplane distance = 400 nm, astigmatism value = 1.4 λ/2π. The region of interest (ROI) size was set to 32 × 32 pixels.

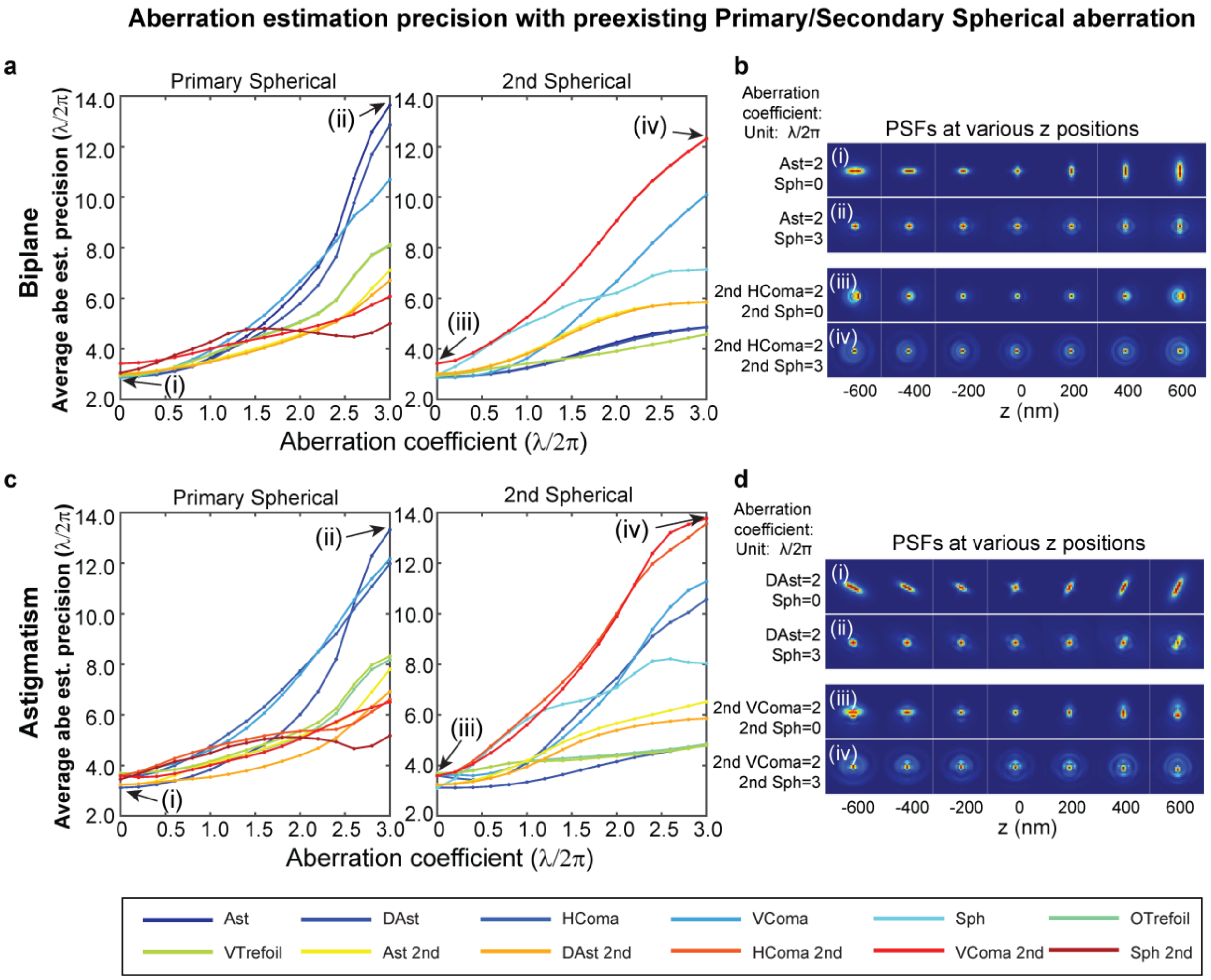

**Supplementary Fig. 20: Aberration estimation precision with preexisting primary and secondary spherical aberrations.** (a, c) Precision of aberration estimation when estimating each aberration at an amplitude of 2 λ/2π while introducing varying levels of primary (left) and secondary (right) spherical in (a) biplane and (c) astigmatism-based system. The precision values are calculated by averaging within the -600 nm to 600 nm z range. The rainbow colors represent different aberration modes, ranging from vertical astigmatism to secondary spherical aberration. (b, d) Examples of PSFs within the -600 nm to 600 nm range, corresponding to cases as indicated in the figures. Simulation conditions: $\lambda$ = 680 nm, $I$ = 0.5 photons/plane for the biplane system, $I$ = 1 photon for the astigmatism system, $bg$ = 0, $NA$ = 1.4, $n_{obj}$ = 1.52, pixel size = 130 nm, biplane distance = 400 nm, astigmatism value = 1.4 λ/2π. The region of interest (ROI) size was set to 32 × 32 pixels.

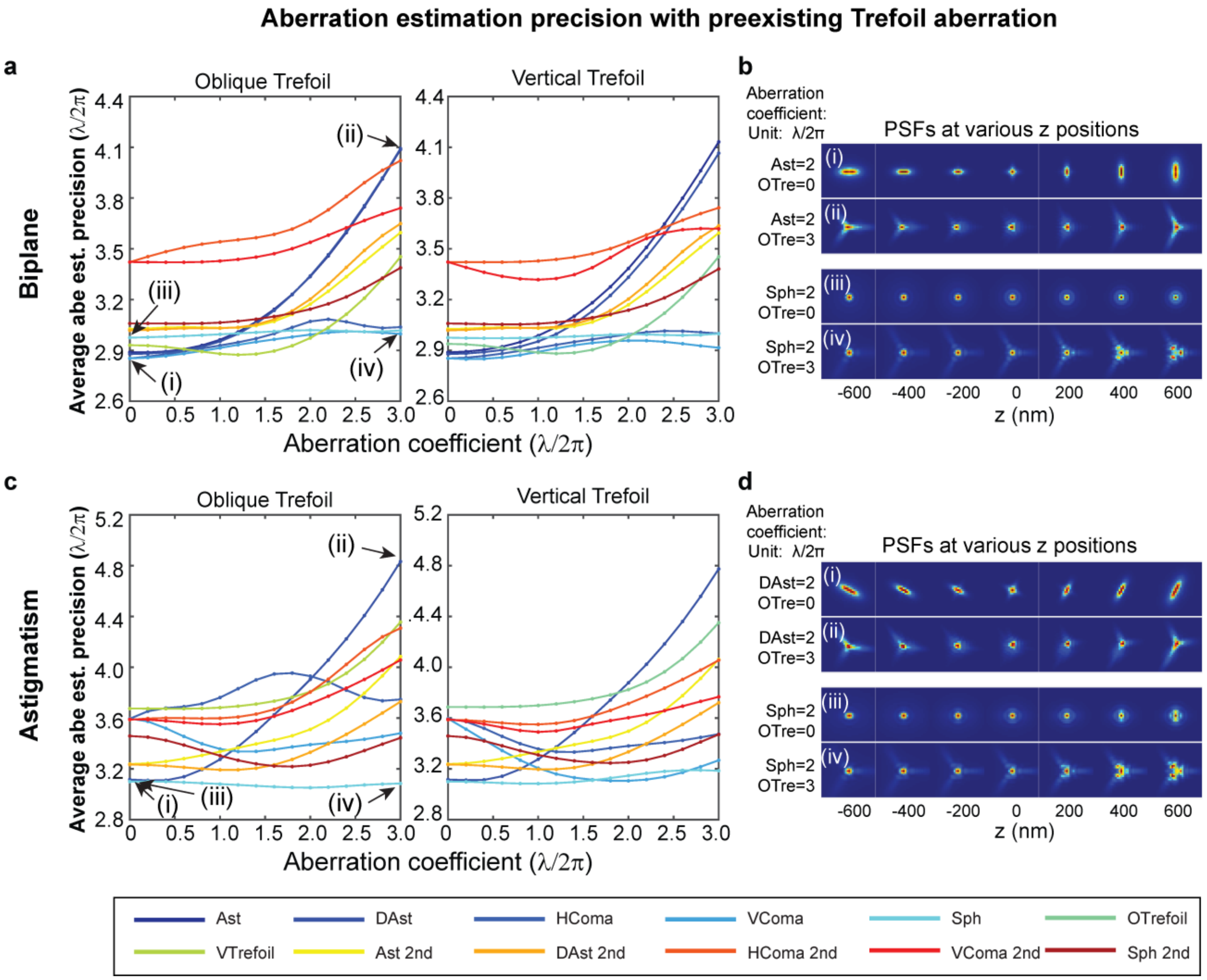

**Supplementary Fig. 21: Aberration estimation precision with preexisting trefoil aberrations.** (a, c) Precision of aberration estimation when estimating each aberration at an amplitude of 2 λ/2π while introducing varying levels of oblique (left) and vertical (right) trefoil in (a) biplane and (c) astigmatism system. The precision values are calculated by averaging within the -600 nm to 600 nm z range. The rainbow colors represent different aberration modes, ranging from vertical astigmatism to secondary spherical aberration. (b, d) Examples of PSFs within the -600 nm to 600 nm range, corresponding to cases as indicated in the figures. Simulation conditions: $\lambda$ = 680 nm, $I$ = 0.5 photons/plane for the biplane system, $I$ = 1 photon for the astigmatism system, $bg$ = 0, $NA$ = 1.4, $n_{obj}$ = 1.52, pixel size = 130 nm, biplane distance = 400 nm, astigmatism value = 1.4 λ/2π. The region of interest (ROI) size was set to 32 × 32 pixels.

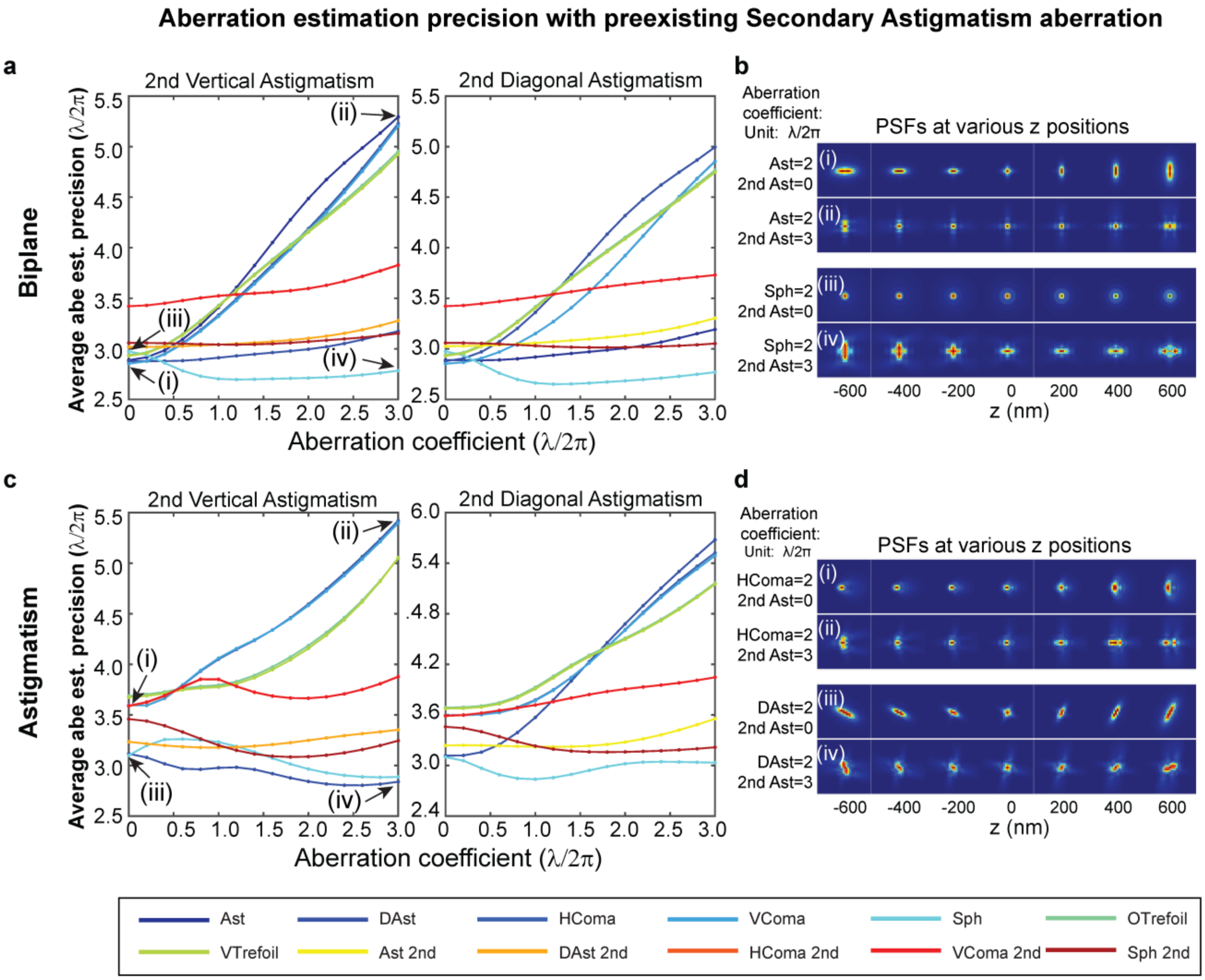

**Supplementary Fig. 22: Aberration estimation precision with preexisting secondary astigmatism aberrations.** (a, c) Precision of aberration estimation when estimating each aberration at an amplitude of 2 λ/2π while introducing varying levels of secondary vertical (left) and diagonal (right) astigmatism in (a) biplane and (c) astigmatism system. The precision values are calculated by averaging within the -600 nm to 600 nm z range. The rainbow colors represent different aberration modes, ranging from vertical astigmatism to secondary spherical aberration. (b, d) Examples of PSFs within the -600 nm to 600 nm range, corresponding to cases as indicated in the figures. Simulation conditions: $\lambda$ = 680 nm, $I$ = 0.5 photons/plane for the biplane system, $I$ = 1 photon for the astigmatism system, $bg$ = 0, $NA$ = 1.4, $n_{obj}$ = 1.52, pixel size = 130 nm, biplane distance = 400 nm, astigmatism value = 1.4 λ/2π. The region of interest (ROI) size was set to 32 × 32 pixels.

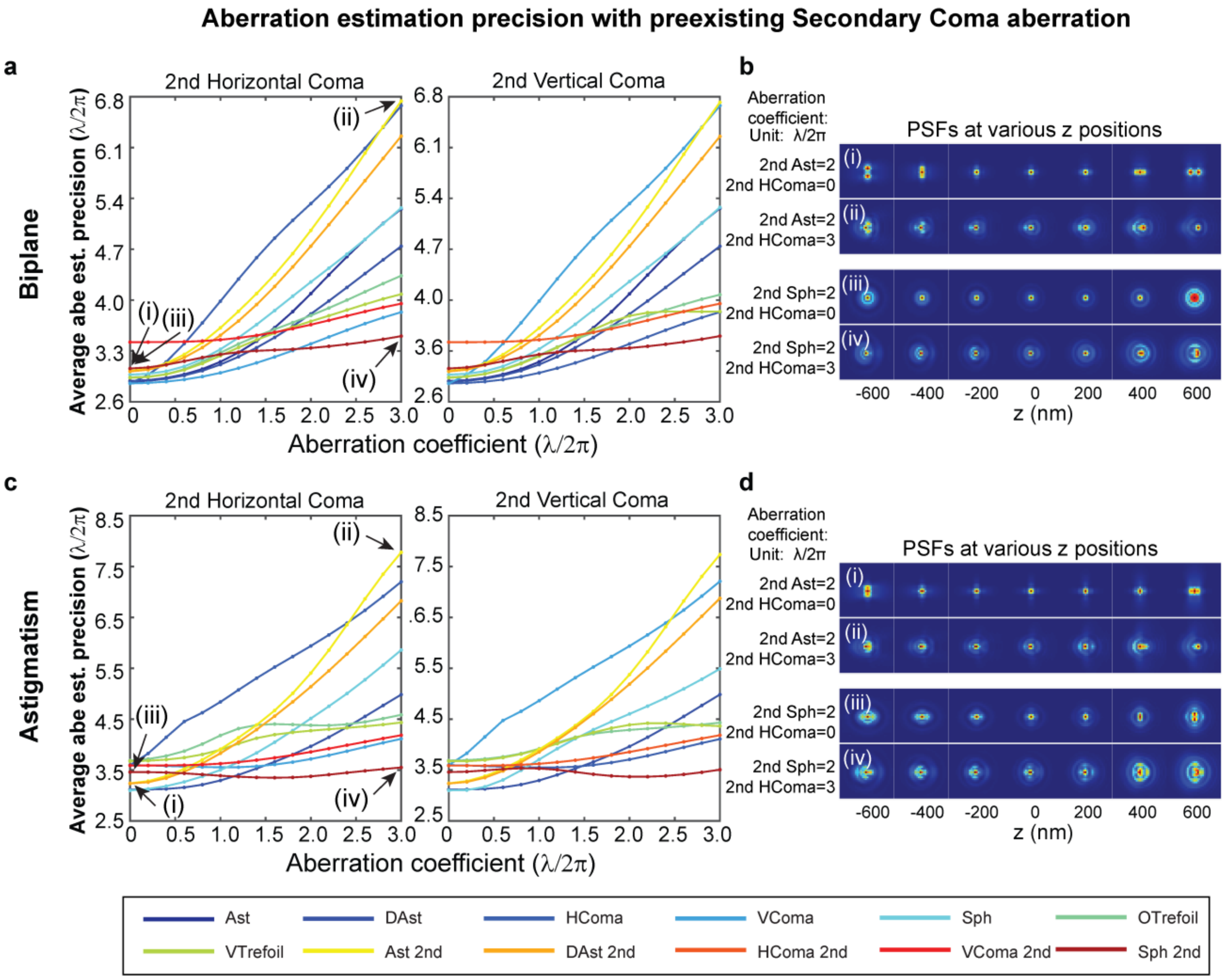

**Supplementary Fig. 23: Aberration estimation precision with preexisting secondary coma aberrations.** (a, c) Precision of aberration estimation when estimating each aberration at an amplitude of 2 $\lambda/2\pi$ while introducing varying levels of secondary horizontal (left) and vertical (right) coma in (a) biplane and (c) astigmatism system. The precision values are calculated by averaging within the -600 nm to 600 nm z range. The rainbow colors represent different aberration modes, ranging from vertical astigmatism to secondary spherical aberration. (b, d) Examples of PSFs within the -600 nm to 600 nm range, corresponding to cases as indicated in the figures. Simulation conditions: $\lambda$ = 680 nm, $I$ = 0.5 photons/plane for the biplane system, $I$ = 1 photon for the astigmatism system, $bg$ = 0, $NA$ = 1.4, $n_{obj}$ = 1.52, pixel size = 130 nm, biplane distance = 400 nm, astigmatism value = 1.4 $\lambda/2\pi$. The region of interest (ROI) size was set to 32 × 32 pixels.

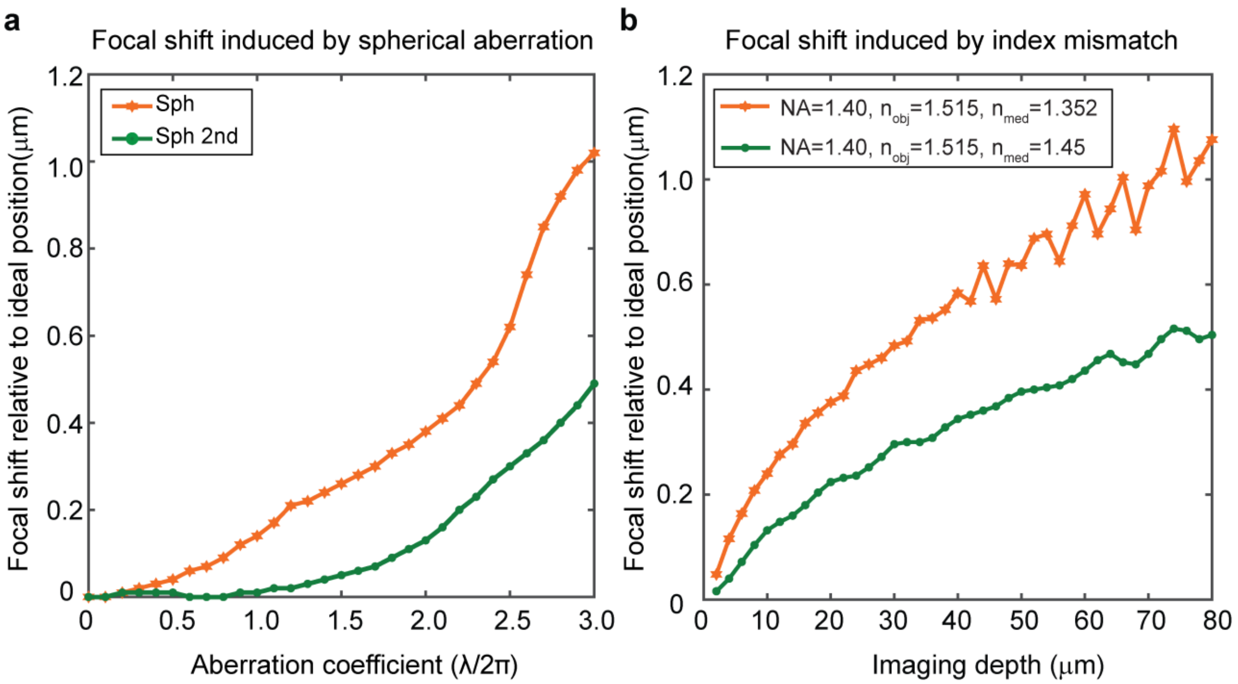

**Supplementary Fig. 24: Focal shift induced by spherical, secondary spherical and refractive index mismatch aberration.** (a) Axial focal shift caused by different amounts of primary and secondary spherical aberration. Simulation conditions: $\lambda$ = 680 nm, $I$ = 1 photon, $bg$ = 0, $NA$ = 1.4, $n_{obj}$ = 1.52, pixel size = 130 nm. (b) Axial focal shift caused by refractive index mismatch aberration at different imaging depths. Simulation conditions: $\lambda$ = 680 nm, $I$ = 1000 photons, $bg$ = 10, $NA$ = 1.4, $n_{obj}$ = 1.52, pixel size = 130 nm.
